# Probabilistic tensor decomposition extracts better latent embeddings from single-cell multiomic data

**DOI:** 10.1101/2022.08.26.505382

**Authors:** Ruohan Wang, Jianping Wang, Shuai Cheng Li

## Abstract

Single-cell sequencing technology enables the simultaneous capture of multiomic data from multiple cells. The captured data can be represented by tensors, i.e., the higher-rank matrices. However, the proposed analysis tools often take the data as a collection of two-order matrices, renouncing the correspondences among the features. Consequently, we propose a probabilistic tensor decomposition framework, SCOIT, to extract embeddings from single-cell multiomic data. To deal with sparse, noisy, and heterogeneous single-cell data, we incorporate various distributions in SCOIT, including Gaussian, Poisson, and negative binomial distributions. Our framework can decompose a multiomic tensor into a cell embedding matrix, a gene embedding matrix, and an omic embedding matrix, allowing for various downstream analyses. We applied SCOIT to seven single-cell multiomic datasets from different sequencing protocols. With cell embeddings, SCOIT achieves superior performance for cell clustering compared to seven state-of-the-art tools under various metrics, demonstrating its ability to dissect cellular heterogeneity. With the gene embeddings, SCOIT enables cross-omics gene expression analysis and integrative gene regulatory network study. Furthermore, the embeddings allow cross-omics imputation simultaneously, outperforming conventional imputation methods with the Pearson correlation coefficient increased by 0.03-0.28.

## Introduction

Single-cell technologies enable cellular heterogeneity dissection at high resolution. Starting with single-cell RNA sequencing (scRNA-seq)^1^, the technologies have been extended to DNA methylation, proteomics, and chromatin accessibility, providing unprecedented opportunities to study biological systems from various perspectives^2^. More recently, technological advances have allowed the capture of different types of molecules and epigenetic data from the same cell^3,4^. Multiomic data capture information from multiple sources at a single cell resolution, allowing more comprehensive studies of cellular heterogeneity and biological systems^5,6^. Cao *et al*. simultaneously measured the expression of RNA and chromatin accessibility of mouse kidney cells. They inferred the target genes for distal *cis*-regulatory elements from the covariance of the two arrays^7^. Yan *et al*. showed that the joint profile of DNA methylation, RNA expression, and chromatin accessibility of human and mouse oocytes revealed the evolution of gene body methylation^8^. Luo *et al*. discovered the relations between DNA methylation and gene expression for neuronal cells with multiomic measurement^9^.

Despite advances in sequencing technology^7–14^, the integrated analysis of single-cell multiomic data is statistically challenging. The challenges come from two aspects. First, single-cell data are inherently sparse. Zero-count occurrences can be due to either actual absence or technical errors, i.e., dropout events, in the scRNA-seq data^15^. The problem is more severe for the chromatin accessibility data^16,17^. Second, single-cell sequencing data exhibit high noise, and the high noise is due to low capture efficiency and shallow sequencing depth^18,19^, leading to a deviated representation of the actual distribution.

The heterogeneity of multiomics poses another challenge for consensus inferences^4^. First, sequencing data from different omics arise from distinct biological variations and technical bias^20^. Second, the data to be integrated have various types. RNA expression and proteomic data can be discrete or continuous, whereas epigenomic data are binary. The distinct features of the omic data impede statistical modeling.

Methods developed for single-cell multiomic data integration are emerging. Matcher^21^ applies manifold alignment to learn the low-dimensional representation of cells and project them into a shared space. UnionCom^22^ advances the manifold alignment algorithm to achieve soft alignment between cells in all omics. MOFA^23^ and the updated version MOFA+^24^ perform joint matrix factorization to infer the cell representation in a latent space. scAI^17^ extends the matrix factorization formulation by aggregating neighbors’ signals to rectify the sparsity of the data. totalVI^20^ leverages variational autoencoder to learn the representations of cells in omics. The latest version of Seurat^25^ applies the weighted nearest neighbor to build a joint cell-cell graph for downstream analysis.

The aforementioned methods are built on jointly projecting the cells into a latent space or graph, while ignoring the integration at the gene level. The main reason is that most of these methods represent the multiomic data as a feature-wise concatenated matrix. In this context, cells are embedded in a shared space, whereas genes are embedded in separated spaces for multiple omics. Arranging the multiomic data in a two-dimensional matrix fails to fully utilize the integrated information. Tensor, a higher-rank generalization of a matrix, offers a natural representation of data with multiple facets. A tensor can order the variables along different tensor dimensions, including cell, feature, and omic. Then, we can derive the integrated information for the variables by decomposing and embedding the tensor.

In this work, we propose a probabilistic tensor decomposition framework to extract embeddings from single-cell multiomic data. To deal with sparse, noisy, and heterogeneous single-cell data, we applied various distributions, including the Gaussian distribution, Poisson distribution, and negative binomial distribution, to model different data types. The framework can decompose a multiomic tensor into a cell embedding matrix, a gene embedding matrix, and an omic embedding matrix, allowing for various downstream analyses. We implemented the framework in an open source package named SCOIT (**S**ingle **C**ell multi-**O**mics data **I**ntegration with **T**ensor decomposition).

We applied SCOIT to seven single-cell multiomic datasets from various sequencing protocols, covering DNA methylation data, RNA expression data, proteomics data, and chromatin accessibility data. First, with integrated cell embeddings, SCOIT achieved better clustering accuracy than the seven state-of-the-art methods on heterogeneous datasets. Furthermore, the integrated gene embeddings allowed us to study gene expressions at different levels and investigate the post-transcriptional gene regulatory network, which single-omic data cannot offer. Moreover, we demonstrate that the embeddings from SCOIT allow multiomic imputation, outperforming the conventional imputation tools measured by Pearson correlation coefficients and root mean square error. SCOIT provides a versatile framework that can be flexible to new downstream analyses and applications.

## Results

### SCOIT overview

This study proposes and implements SCOIT, a probabilistic tensor decomposition framework for integrating single-cell multiomic data. SCOIT accepts datasets from multiple omics, with missing values allowed, and it processes the data in two steps (Figure 1 A). First, it constructs a multiomic tensor with a union set of features. Second, it performs the probabilistic tensor decomposition with a user-specified distribution. SCOIT generates embedding matrices for omics, cells, and genes.SCOIT incorporates the global and local embeddings to capture global and local variability. The embeddings enable downstream analyses, including cell clustering, cell-cell graph construction, gene expression study, data imputation, *etc*. (Figure 1 B). SCOIT is available at https://github.com/deepomicslab/SCOIT.

**Figure 1.**
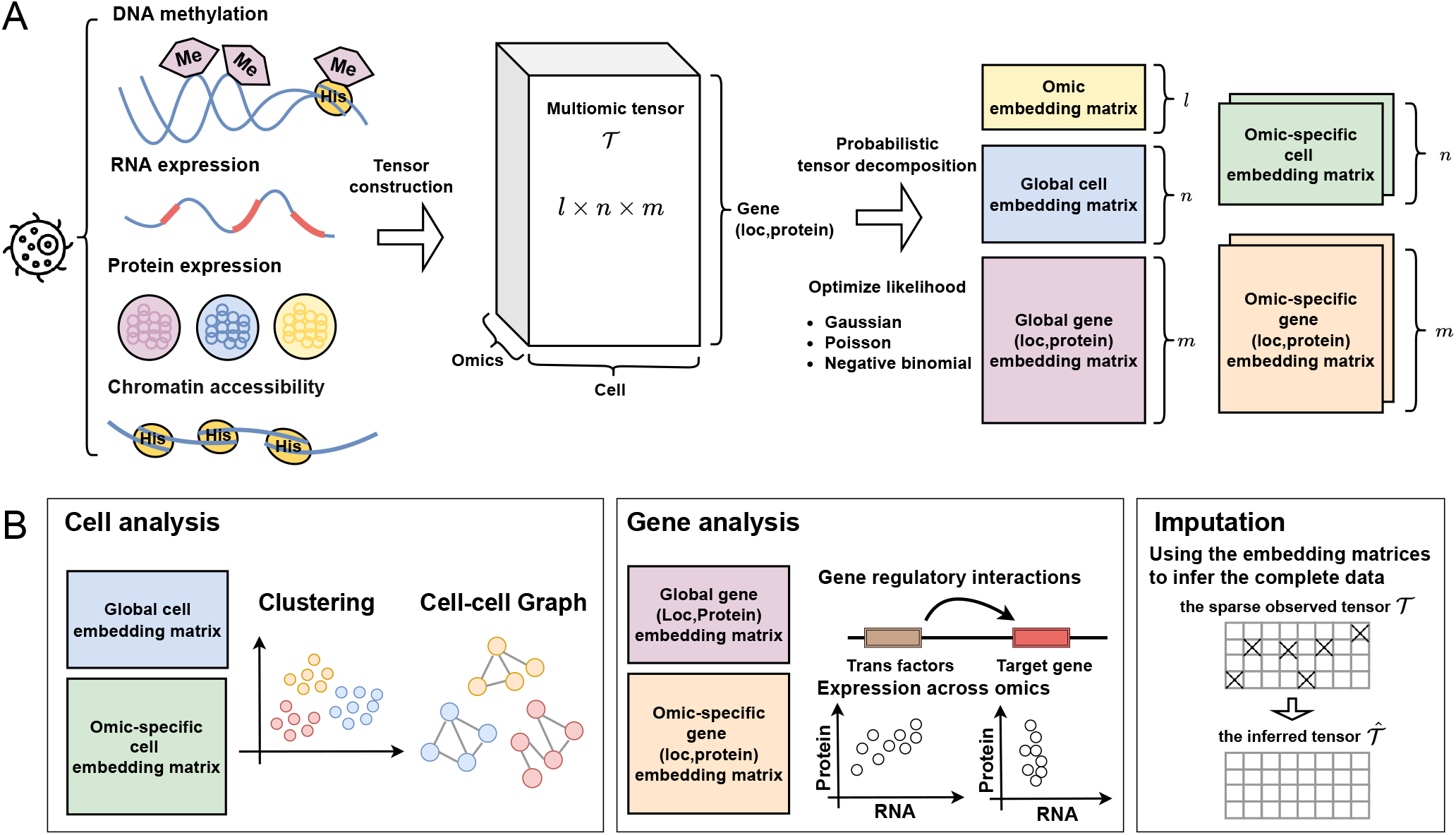
SCOIT provides a probabilistic tensor decomposition framework for the integration of single-cell multiomic data. **A. Overview:** SCOIT constructs a multiomic tensor with the input of single-cell sequencing data from multiple omics. Then following a user-defined distribution, SCOIT decomposes the tensor into an omic embedding matrix, a global cell embedding matrix, a global gene (location, protein) embedding matrix, an omic-specific cell embedding matrix, and an omic-specific gene (location, protein) embedding matrix. **B. Downstream analysis:** the cell embedding matrices can be applied to cell clustering analysis and cell nearest neighbor (NN) graph construction; the gene embedding matrices can be applied to study the gene regulatory interactions and the expression across multiomics; the inferred tensor, computed from the embedding matrices, can be used for imputation.

### SCOIT integrates RNA expression and DNA methylation data from sc-GEM

Sc-GEM jointly profiles RNA expression and DNA methylation at single-cell level^10^. We applied SCOIT to analyze 224 human fibroblast cells in various stages of the reprogramming process sequenced with sc-GEM. Cells are labeled by their reprogramming stages. The RNA expression data contain 34 genes, and the DNA methylation data includes 27 genes. We construct the multiomic tensor with the union set of the genes.

SCOIT generates a global cell embedding matrix that gives each cell an integrated representation. According to the clustering of *k*-means with the cell embeddings by SCOIT and the benchmark methods, the SCOIT embedding achieves the best clustering performance according to various metrics (Figure 2 A), with scAI ranked the second. We visualize the cell embeddings with uniform manifold approximation and projection (UMAP). As shown in Figure 2 B, the projection of the SCOIT cell embeddings presents a linear structure that shares the same order as the trajectory of the reprogramming process. We also provide the projection of the embeddings learned by the benchmark methods in Supplementary Figure S1 for comparison. Furthermore, we analyze the RNA expression and DNA methylation patterns using the global gene embedding matrix.

**Figure 2.**
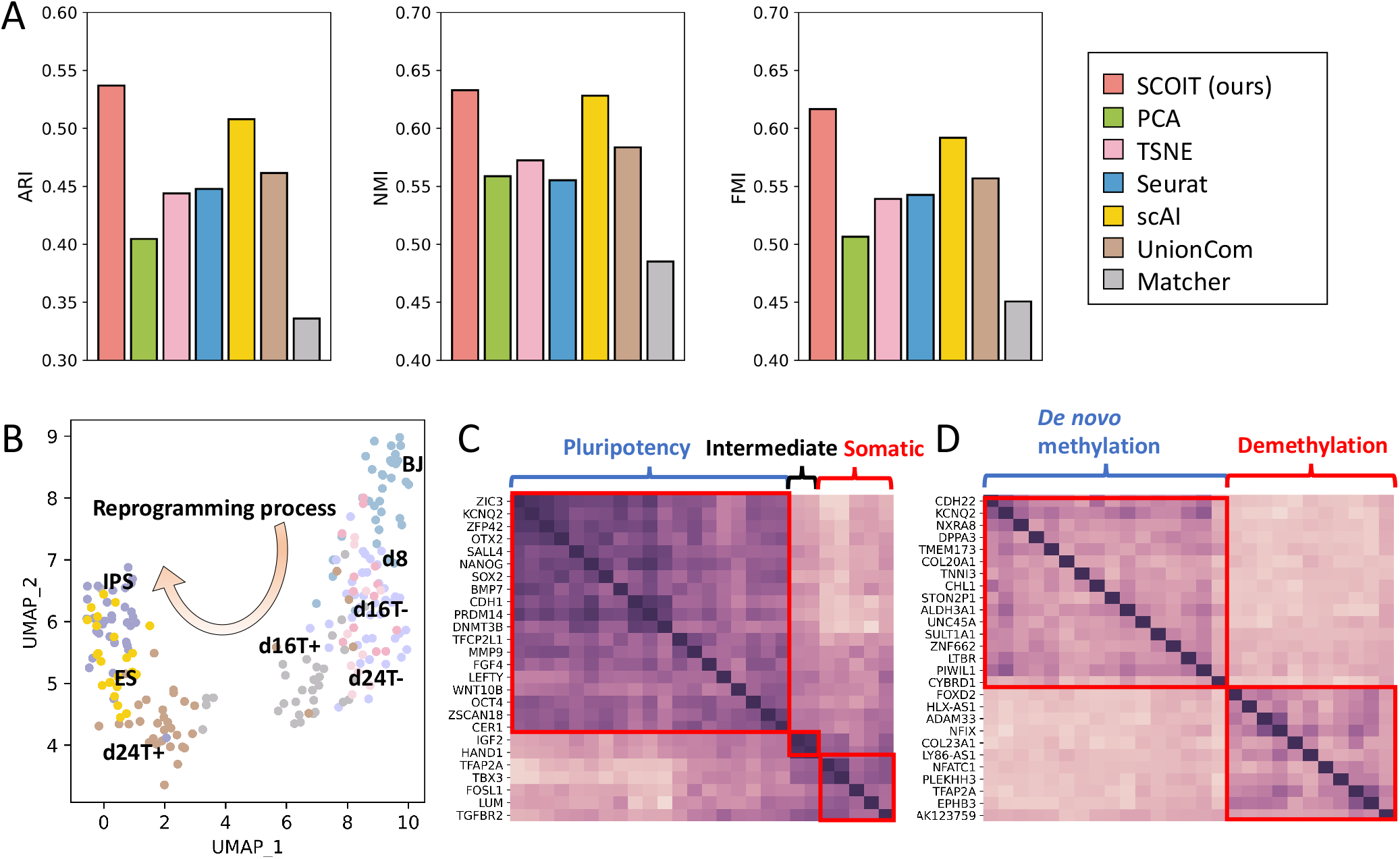
SCOIT reveals cellular heterogeneity and gene expression or methylation patterns from integrating gene expression and DNA methylation data of human fibroblast cells. **A**. Comparison of *k*-means clustering performance with the input of cell embeddings generated by different methods, measured by ARI, NMI, and FMI. The higher bar corresponds to more concordance between the predicted and true labels. **B**. The UMAP projections of the cell embeddings generated by SCOIT. Each point represents a cell color-coded by the true label. The arrow shows the time-ordered reprogramming process. **C. and D**. The heatmap of correlations between the gene embeddings. Darker color indicates a higher Pearson correlation coefficient. The genes are grouped in red rectangles according to the gene expression pattern (C) and DNA methylation pattern (D).

In the previous study^10^, the genes in the RNA expression dataset are classified into pluripotent, intermediate, and somatic groups according to their transcriptional changes during the reprogramming process. Genes in the DNA methylation dataset are grouped into *de novo* methylation and demethylation according to their dynamic methylation patterns. Using the global gene embedding matrix, we computed the Pearson correlation coefficients between the genes. Figure 2 C and D show that the correlations for genes in the same group are higher, indicating that global gene embeddings capture shared patterns of RNA expression and DNA methylation.

### SCOIT integrates RNA expression and chromatin accessibility data

We consider three datasets to evaluate the ability of SCOIT to integrate RNA expression and chromatin accessibility data. They are 8,837 adult mouse kidney cells sequenced with sci-CAR^7^, 5,081 neonatal mouse cerebral cortex cells sequenced with SNARE-seq^11^, and 10,309 adult mouse cerebral cortex cells sequenced with SNARE-seq^11^. We independently perform dimensional reduction (see *Omics data processing* subsection) on the RNA expression and chromatin accessibility data to reduce the data size. SCOIT constructs the tensor with the processed matrices.

With the global cell embedding matrix generated by SCOIT, we construct a K-Nearest Neighbor (KNN) graph and a Shared Nearest Neighbor (SNN) graph (see *Cell clustering and graph construction* subsection). Then we apply Leiden^26^ to perform community detection for the two graphs. We also used the cell embeddings of the benchmark methods to construct graphs for comparison. SCOIT outperforms other methods significantly in community detection with both the KNN graph and the SNN graph for the three datasets (Figure 3 A and C, Supplementary Figure S2 A). The UMAP visualizations of the global cell embedding matrix (Figure 3 B and D, Supplementary Figure S2 B) show that cells with the same types cluster together and cells with different types are separated. Supplementary Figures S3-S5 show the UMAP visualization of the embeddings provided by the benchmark methods. The results suggest that the global cell embeddings learned by SCOIT give good representations of the cells.

**Figure 3.**
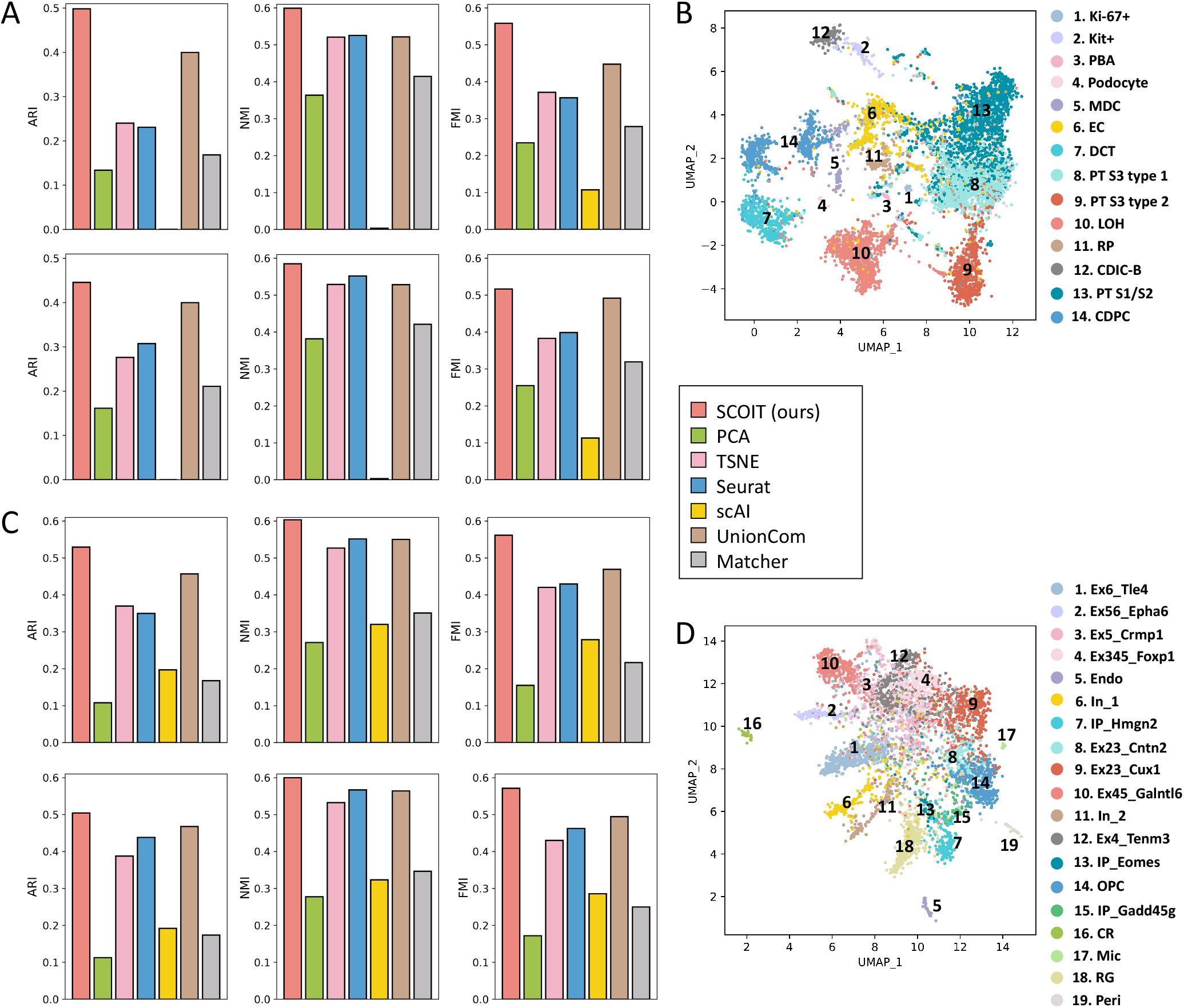
SCOIT reveals cellular heterogeneity from integrating RNA expression and chromatin accessibility data. **A. and C**. For data from adult mouse kidney cells sequenced by sci-CAR (A) and data from neonatal mouse cerebral cortices cells sequenced by SNARE-seq (C), we compare the Leiden community detection performance for the KNN graph (top) and SNN graph (bottom), constructed with cell embeddings generated by different methods. ARI, NMI, and FMI measure the performance. **B. and D**. For data from adult mouse kidney cells sequenced by sci-CAR (B) and data from neonatal mouse cerebral cortices cells sequenced by SNARE-seq (D), we show the UMAP projections of the cell embeddings generated by SCOIT. Each point represents a cell color-coded by the true label. The cell types are shown on the right of the scatter plot.

### SCOIT integrates RNA expression and proteomics data from PEA/STA

PEA/STA^12^ provides a dataset containing RNA expression and proteomic data sequenced from 210 human glioblastoma cells to investigate the treatment effect of BMP4. Eighty-eight genes are measured in the RNA expression dataset, and 78 genes are measured in the proteomic dataset. A previous study^12^ shows a low correlation between RNA expression and protein expression at the single-cell level in this dataset. Therefore, we increase the coefficient of the penalty term for global gene embeddings, forcing the model to learn omic-specific gene embeddings (see *Model regularization* subsection).

We apply *k*-means clustering to global cell embeddings to distinguish cells with and without BMP4 treatment. SCOIT embeddings achieve the best clustering performance among all benchmark methods (Figure 4 A). UnionCom, which relies on a similar topological structure alignment, performs the worst for this dataset, probably due to the low correlation between the two omics. The UMAP visualization for cell embeddings also shows a more apparent separation between control and BMP4-treated cells than from the benchmark methods (Figure 4 B and Supplementary Figure S6).

**Figure 4.**
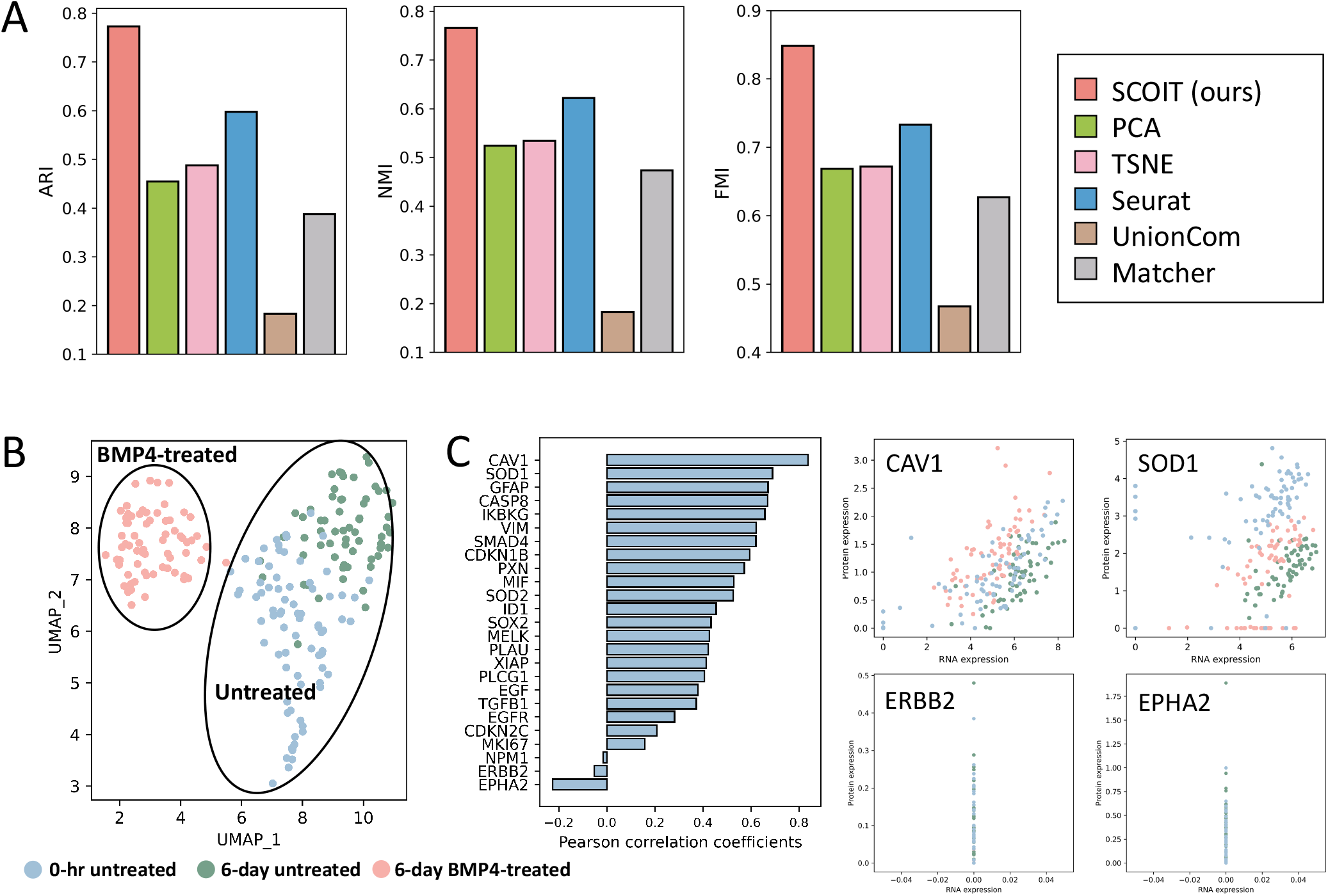
SCOIT reveals cellular heterogeneity and the correlation between RNA and protein expression levels from integrating RNA expression and proteomics data of human breast adenocarcinoma cells. **A**. Comparison of *k*-means clustering performance with the input of cell embeddings generated by different methods, measured by ARI, NMI, and FMI. **B**. The UMAP projections of the cell embeddings generated by SCOIT. Each point represents a cell color-coded by the true label. The circles separate the untreated and treated cells. **C**. The left bar plot shows the correlation between gene embeddings from RNA expression and proteomics generated by SCOIT. The higher bar corresponds to a higher Pearson correlation coefficient. The right scatter plots show the RNA and protein expression levels of CAV1, SOD1, ERBB2, and EPHA2, for all cells. The points are color-coded by the true label, sharing the legend in B.

Next, we study the correlations between RNA and protein expression levels. We compute the Pearson correlation coefficients between the RNA-specific and protein-specific gene embeddings. The distribution graph shows a lower correlation between the expression levels of RNA and proteins for most genes (Supplementary Figure S7), which is consistent with the previous report^12^. We present the Pearson correlation coefficients for the genes measured in both omics in the bar plot in Figure 4 C. We show the expression levels of RNA and proteins for the two genes with the highest correlations (CAV1 and SOD1) and the two with the least correlations (ERBB2 and EPHA2). CAV1 and SOD1 demonstrate strong correlations, while ERBB2 and EPHA2 do not, consistent with the Pearson correlation coefficients calculated from gene embeddings.

### SCOIT integrates RNA expression and epitope data from CITE-seq

CITE-seq^13^ simultaneously measures RNA expression and epitope data for single cells. We apply SCOIT to a CITE-seq dataset with 8,617 cord blood mononuclear cells. The RNA expression dataset contains 36,280 genes, and the epitope dataset contains 13 surface proteins. SCOIT constructs a multiple-feature tensor (Supplementary methods S1.1) with the two datasets.

We conduct *k*-means clustering on the cell embeddings generated by SCOIT and the benchmark methods. With the labels identified by the protein markers as the ground truth, SCOIT and Seurat achieve superior performance according to various metrics (Figure 5 A). The epitope data contain more information to determine cell types. Seurat automatically determines the importance of each omics and weights according to the importance during integration. PCA and TSNE have the poorest performance as they treat the features of each omics equally. UMAP visualizations for cell embeddings are shown in Supplementary Figure S8.

**Figure 5.**
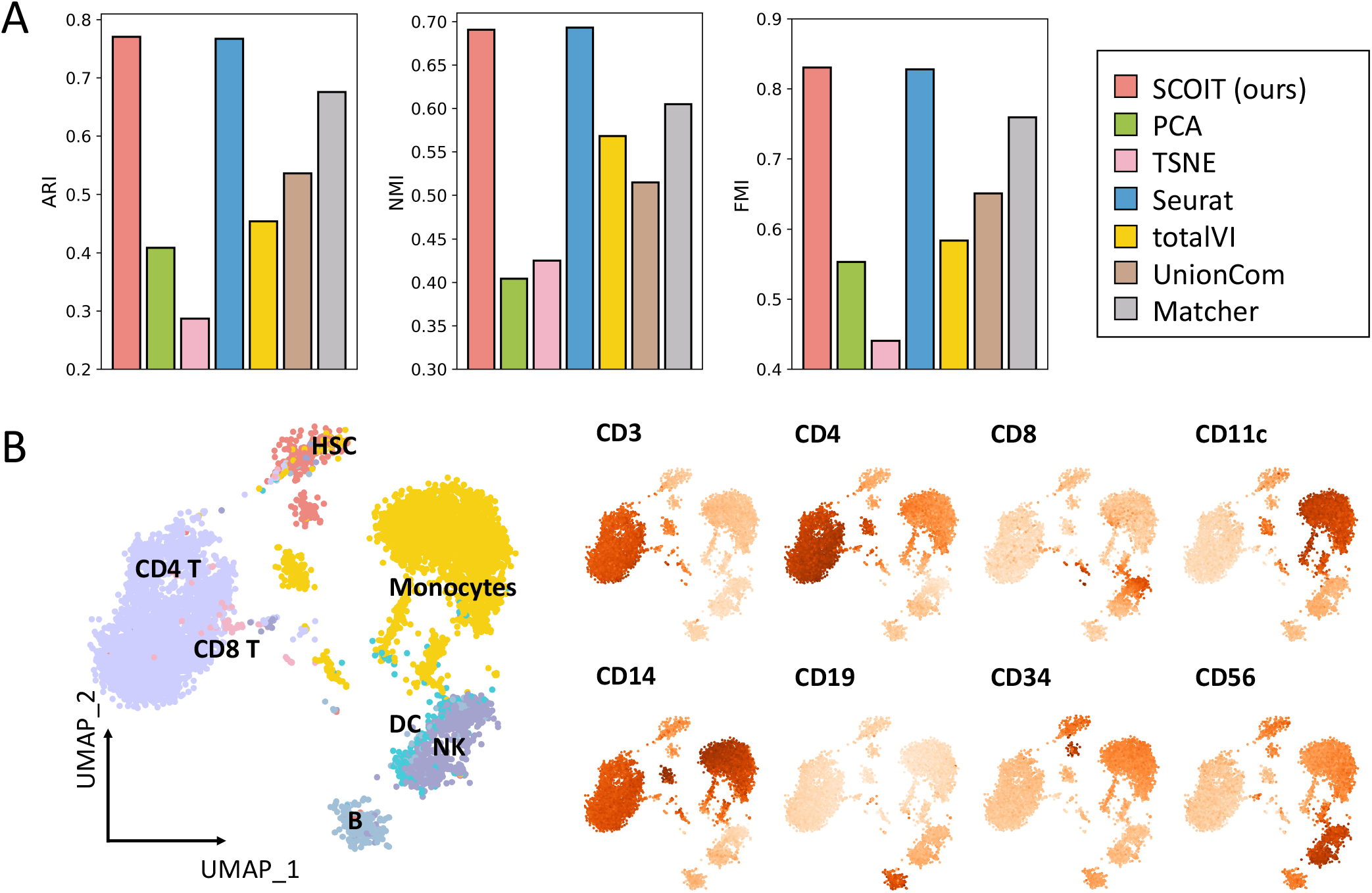
SCOIT reveals cellular heterogeneity and identifies distinct antigens from integrating RNA expression and epitopes data of cord blood mononuclear cells. **A**. Comparison of *k*-means clustering performance with the input of cell embeddings generated by different methods, measured by ARI, NMI, and FMI. **B**. The left scatter plot shows the UMAP projections of the ADT signals. Each point represents a cell color-coded by the true label. The right scatter plots show the projections of the SCOIT-generated epitope embeddings on cells. Darker color indicates a higher value.

We then investigated the correlations between the RNA and protein expression levels for the CITE-seq data. We identified six genes that encode the corresponding epitope and calculated the Pearson correlation coefficients between the gene embeddings and the 13 epitope embeddings generated by SCOIT. As shown in Supplementary Figure S9, gene embeddings have the highest coefficients with their encoded epitope embeddings, indicating a strong correlation between RNA and protein expression levels.

Moreover, we applied epitope embeddings to study epitope distributions of different cell types. We identified eight cell-type-specific epitopes and projected their embeddings onto the global cell embeddings (by an inner product). As shown in Figure 5 B, embedding projection results in a higher value for cells with epitope expression. The result indicates that the embeddings capture the heterogeneity of epitopes.

### SCOIT integrates SCoPE2 proteomic data and 10X RNA expression data

SCoPE2^14^ provides a proteomic dataset with 3,042 proteins in 1,490 single monocytes and macrophages and parallel measurement of RNA expression by 10× Genomics. Cells in the two datasets have been paired with the first shared principal components^14^. Based on the set of genes from the two datasets that are combined, we construct a multiomic tensor as the input of SCOIT.

Applying the *k*-means method to the cell embeddings generated by SCOIT and the benchmark methods again suggests that the SCOIT embedding outperforms all other embeddings (Figure 6 A). The UMAP projections of the global cell embeddings given by SCOIT present a better collection of cell types (Figure 6 B and Supplementary Figure S10), suggesting that SCOIT generates more informative representations for cells.

**Figure 6.**
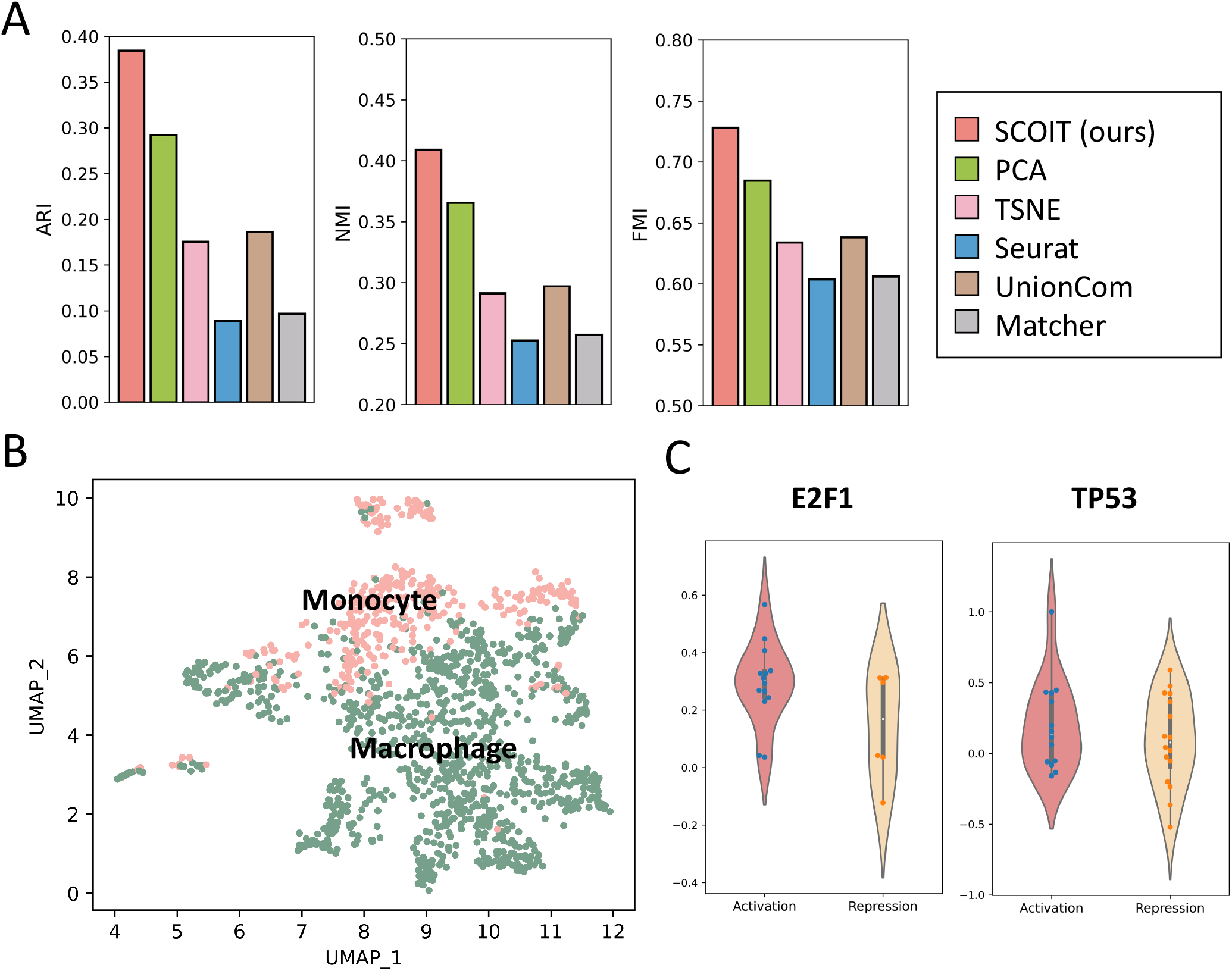
SCOIT reveals cellular heterogeneity and dissects regulatory interactions by integrating RNA expression and proteomics data of innate immune cells. **A**. Comparison of *k*-means clustering performance with the input of cell embeddings generated by different methods, measured by ARI, NMI, and FMI. **B**. The UMAP projections of cell embeddings generated by SCOIT. Each point represents a cell color-coded by the true label. **C**. For transcription factors E2F1 and TP53, the violin plots show the correlation distributions between the gene embedding and its target gene embeddings (left for activated genes and right for repressed genes), generated by SCOIT. The dots inside the boxes show the Pearson correlation coefficients.

In addition, we apply global gene embeddings to study post-transcriptional gene regulation. We use TRRUST^27^, a reference database for the human gene regulatory network, to identify target genes (including activated genes and repressed genes) for transcription factors (TF) in the SCoPE2 dataset. TFs with more than five activated genes and repressed genes in the SCoPE2 dataset are used for analysis to ensure statistical significance, resulting in two TFs, E2F1, and TP53 (Supplementary Table S1). Then we compute the Pearson correlation coefficients between the TF embeddings and their target gene embeddings to infer the correlations. As expected, TF embeddings correlate positively with activated gene embeddings and negatively with repressed gene embeddings (Figure 6 C). The results also agree with previous findings^14^.

### SCOIT persists its performance with highly sparse and noisy data

To further assess the performance of SCOIT on highly sparse and noisy datasets, we add missing values and Gaussian noises to the sc-GEM dataset. We simulate the *in silico* sparse and noisy datasets under four settings: (1) setting 1% − 30% of the observations in the RNA expression dataset to the missing values; (2) adding Gaussian noise (mean:5, variance:1) to 1% − 30% of the observations in the RNA expression dataset; (3) setting 1% − 30% of the observations in the DNA methylation dataset to the missing values; (4) adding Gaussian noise (mean:1, variance:0.5) to 1% − 30% of the observations in the DNA methylation dataset.

We applied SCOIT to the simulated datasets and performed *k*-means method on the cell embeddings. The results are shown in Supplementary Figure S11. The performance remains relatively stable for the datasets of various missing value distributions and noise amplitudes, with ARI ranging from 0.4857 to 0.5367, NMI ranging from 0.5912 to 0.6294, and FMI ranging from 0.5759 to 0.6095. The results indicate that SCOIT is robust to sparse and noisy datasets, which makes it suitable for single-cell sequencing data.

### SCOIT effectively imputes multiomic data

The multiomic tensor can be reconstructed by the decomposed matrices. SCOIT can be used to impute multiomic data. We set 10% − 50% of the observations in the sc-GEM dataset as missing values to evaluate the imputation performance. We applied two conventional imputation methods, KNNImputer and SimpleImputer, as benchmark methods. We compute the Pearson correlation coefficients, and the root mean square error between the original and imputed values. As shown in Figure 7, SCOIT achieves the best recovery capacity for all simulated datasets, increasing the Pearson correlation coefficients by 0.03-0.28 and decreasing the root mean square error by 0.01-0.10. KNNImputer has been widely used for multiomic data imputation^14,28,29^. The superior performance of SCOIT makes it an effective alternative.

**Figure 7.**
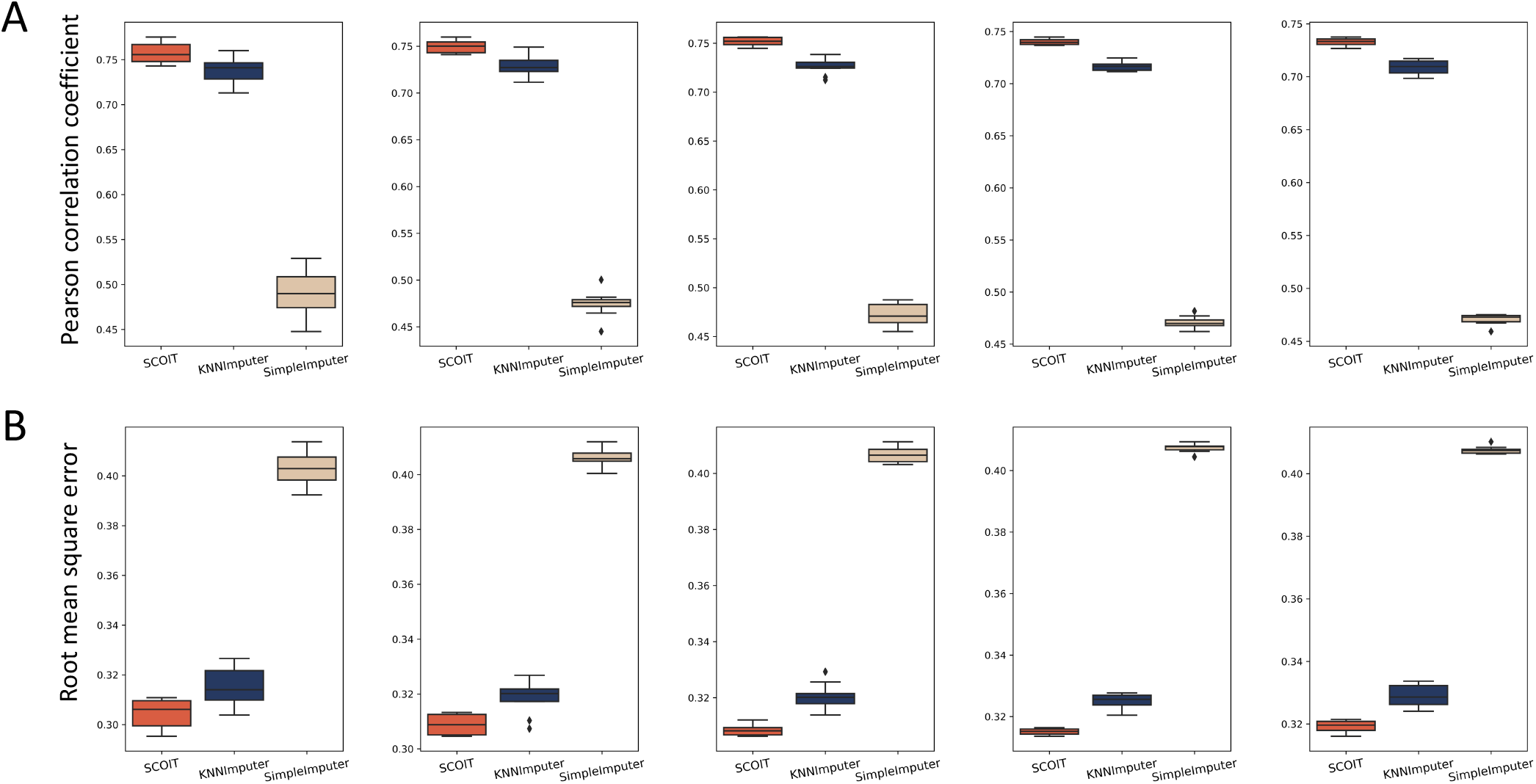
SCOIT achieves the best imputation performance on the sc-GEM dataset compared with KNNImputer and SimpleImputer. From left to right, we set 10%, 20%, 30%, 40%, and 50% of the observations in sc-GEM dataset to missing values (“NA”) and compare the imputation performance concerning **A**. Pearson correlation coefficient and **B**. root mean square error.

## Discussion

Various single-cell multiomics sequencing protocols are emerging. Consequently, there is a growing need for a flexible and scalable method to pool the information from each molecular layer and deliver more comprehensive profiles for complex biological systems. Most existing methods treat each omics data as an independent matrix and conduct joint operations, such as dimension reduction or matrix factorization, on multiple matrices^4^. Such inputs lead to a compromise on information loss. For instance, sc-GEM jointly profiles RNA expression and DNA methylation, with genes as the feature. The matrix operation loses the corresponding gene information. Moreover, representing the multiomic data as a tensor can fully utilize the data, since each variable (omics, cell, and gene) can be modeled as one dimension of the tensor.

We propose a probabilistic tensor decomposition framework to extract information from the multiomic tensor. Compared to the conventional tensor decomposition algorithm, such as canonical polyadic (CP) and Tucker, our framework generates global and local matrices to interpret multilayered biological information. With the adoptive penalty coefficients, users can obtain multiple sources of variation for various downstream analyses. In our case studies, we applied the local gene embeddings to analyze gene expression correlation among omics and the global gene embeddings to study post-transcriptional gene regulations.

Another distinct characteristic of our tensor decomposition framework is the application of various distributions. High noise and sparsity have hindered single-cell sequencing data analysis. A probabilistic model with a suitable distribution can alleviate noise by modeling variations^30^. SCOIT provides various distributions to deal with heterogeneous multiomic datasets. In our experiment, we applied the Gaussian distribution for the continuous data type and the negative binomial distribution for the count data with high variance.

We expect the probabilistic tensor decomposition framework to be helpful beyond multiomic data integration. The framework applies to data with multiple variations. Each underlying source of variation is set as one dimension of the tensor, and the framework provides the embedding for each variable. For instance, our framework corrects the batch effect. Batch-specific systematic variations can be modeled as one dimension in the tensor. Using the tensor decomposition framework to the multi-batch tensor (batch× cell× gene), we obtain the embeddings for the three variables. The terms that include batch embeddings in the decomposition formulation can be removed for batch correction.

## Methods

### Multiomic datasets

We collected seven public single-cell multiomic datasets generated by various sequencing protocols, including sc-GEM^10^, sci-CAR^7^, SNARE-seq^11^, PEA/STA^12^, CITE-seq^13^, and SCoPE2^14^. Sc-GEM provided a dataset (SRA accession SRP077853) sequenced from human fibroblast cells that undergo reprogramming, which contains RNA expression and DNA methylation data. Sci-CAR generated a dataset (GEO accession GSE117089) sequenced from an adult mouse kidney, jointly profiling RNA expression and chromatin accessibility data. SNARE-seq provided two datasets (GEO accession GSE126074) sequenced from neonatal and adult mouse cerebral cortices, respectively, including the RNA expression and chromatin accessibility data. PEA/STA offered a dataset sequenced from human glioblastoma cells containing RNA expression and proteomic data. CITE-seq provides a dataset (GEO accession GSE100866) with cellular indexing of RNA expression and epitopes from cord blood mononuclear cells. SCoPE2 quantified proteomics in innate immune cells (MassIVE ID MSV000083945 and MSV000084660) and provided parallel measurements of RNA expression by 10X Genomics (GEO accession GSE142392). We downloaded the processed data and the identified cell types. CITE-seq did not provide the cell type information, so we labeled the cells with marker proteins (see Supplementary Table S2). We summarize the statistics of the datasets in Supplementary Table S3.

### Omic data processing

We applied various data normalization strategies according to data types. To scale the RNA expression data, we performed min-max normalization for the sc-GEM data. For the data from sci-CAR and SNARE-seq, following the practice introduced in Seurat^25^, we applied scTransform^31^ to the RNA expression data and Signac^32^ to the chromatin accessibility data for data normalization and dimension reduction. For the data from PEA/STA and SCoPE2, we performed a min-max normalization for the RNA expression and proteomic data, since they are given as continuous values. For the RNA expression data from the CITE-seq, we kept the gene detected in at least 10% of the cells and applied log normalization as suggested^13^, then applied min-max normalization to the RNA expression and proteomics data.

### Probabilisitc tensor decomposition framework

#### Notation and definition

We denote the number of omic profiles, the number of cells, and the number of genes as *l, n*, and *m*, respectively. Denote the observed multiomic tensor as 𝒯 ∈ ℝ^*l*×*n*×*m*^, which is constructed with multiple cell × gene data matrices from single omics. Denote the inferred tensor as 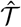, which can be decomposed into the omic embedding matrix **O** ∈ ℝ^*l*×*k*^, the global cell embedding matrix **C** ∈ ℝ^*n*×*k*^, the global gene embedding matrix **G** ∈ ℝ^*m*×*k*^, the omic-specific cell embedding matrix **C**′ ∈ ℝ^*l*×*n*×*k*^, and the omic-specific gene embedding matrix **G**′ ∈ ℝ^*l*×*m*×*k*^, where *k* denotes the rank of decomposition of the tensor.

#### Multiomic tensor construction

Each single-omic data is a two-dimensional matrix (cell × gene (loc)). We can construct a third-order multiomic tensor 𝒯 with the matrices. The data from different omics may contain different genes. We construct the multiomic tensor based on the union set of genes. The features of different omics do not have one-to-one correspondence for some sequencing protocols. For instance, CITE-seq measures RNA expression and epitope data, while one epitope expression is affected by multiple genes. In addition, sci-CAR and SNARE-seq simultaneously profile RNA expression and chromatin accessibility, while the latter gives measurements at different chromatin loci. In these cases, we construct multiple-feature tensors. The decomposition processing for a multiple-feature tensor is analogous to that for one tensor (see Supplementary methods S1.1 and Figure S12). In the following sections, we focus on decomposing single feature (i.e., gene) tensors.

#### Omic-specific tensor decomposition

For an inferred tensor 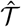, the omic-specific tensor decomposition is expressed element-wisely as

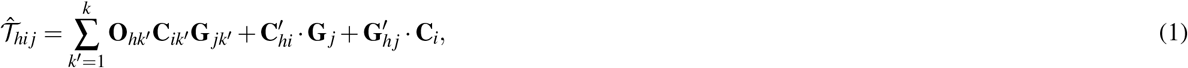

where 1 ≤ *h* ≤ *l*, 1 ≤ *i* ≤ *n*, and 1 ≤ *j* ≤ *m*.

Specifically, the first term is formulated as the sum of the element-wise product for three vectors **O**_*h*_, **C**_*i*_, and **G** _*j*_ ∈ ℝ^*k*^, capturing the global variability for cells and genes across the omics. We adopt a local element-wise product to maintain the variations in each vector (see Supplementary methods S1.2). The second term is the inner product of two vectors 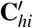 and **G** _*j*_ ∈ ℝ^*k*^, capturing the omic-specific variability for cells. The third term is the inner product of two vectors 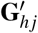 and **C**_*i*_ ∈ ℝ^*k*^, capturing the omic-specific variability of the genes. We give the geometrical interpretation for the formulation of the tensor decomposition in Supplementary Methods S1.3 and Figure S13.

#### Distribution and likelihood objective function

Instead of minimizing the sum-squared distance between 𝒯 and 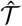, SCOIT adopts likelihood models to resolve the inherent sparsity and high noise of single-cell datasets. SCOIT incorporates various likelihood models that cope with different data types, including a Gaussian distribution model for continuous data, a Poisson distribution model for discrete data, and a negative binomial distribution model for discrete data with high variances.

First, we introduce the likelihood model with Gaussian distribution. The likelihood objective function is as

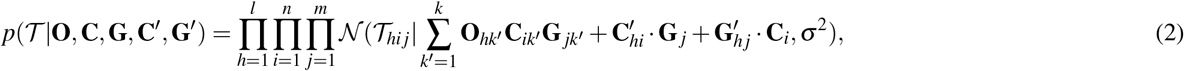

where 𝒩 (*x*|*μ, σ*^2^) is the Gaussian distribution probability density function with mean *μ* and variance *σ* ^2^. *σ* is a hyperparameter in SCOIT.

We recommend the likelihood model with Poisson distribution and negative binomial distribution for the discrete data, i.e., the count data. The likelihood function for the Poisson distribution is as

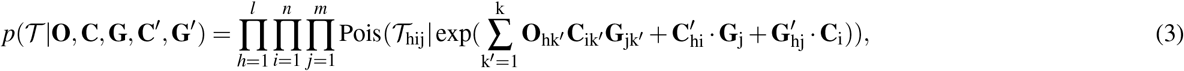

where Pois(x|*λ*) is the probability density function of the Poisson distribution with mean and variance as *λ*.

Since the mean and variance of the Poisson distribution are the same, it is inappropriate for overdispersed data. The negative binomial distribution gives an alternative by placing a gamma prior on *λ* as 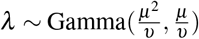, where *μ* is the mean and *υ* is the variance of Gamma distribution. SCOIT applies the constant Fano factor assumption introduced by SAVER^15^, which assumes that the variance scales linearly with the mean under a coefficient. Then the likelihood objective is as

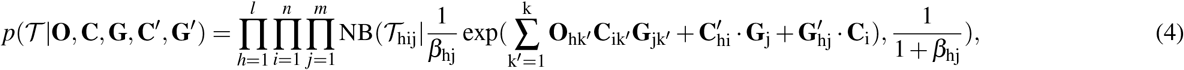

where NB(x|r, p) is the probability density function of the negative binomial distribution with mean 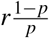 and variance 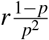 (see Supplementary Methods S1.4 for the reparameterization procedure). *β*_*h j*_ denotes the Fano factor for gene *j* in omics *h*, which is a trainable parameter.

#### Model regularization

SCOIT applies *L*2 regularization to avoid overfitting, increase model interpretability, and control the variability introduced. First, the penalty term shrinks the magnitude of embedding matrix elements and avoids unnecessary model complexity. Second, regularization addresses the sparsity of the embedding matrices, thus encouraging the learning of distinct features for some subsets of genes and samples. Third, the variability of the multiomic data comes from the five embedding matrices, **O, C, G, C**′, and **G**′. We introduce the individually tunable coefficient for the penalty term of each embedding matrix. By tuning the coefficients, we can adjust the integration of information from the global and omic-specific sources. SCOIT also allows for an individual coefficient for each omics to facilitate various downstream analyses.

#### Model optimization

Combining the negative log-likelihood function and the regularization terms, we propose the loss function as

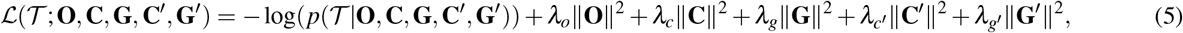

where *λ*_*o*_, *λ*_*c*_, *λ*_*g*_, *λ*_*c*′_, *λ*_*g*′_, are the coefficients for the penalty term of each embedding matrix.

SCOIT performs a gradient descent to minimize the loss function. To be specific, SCOIT initializes **O, C, G, C**′, and **G**′ with a Gaussian distribution (*μ* = 0, *σ* = 0.1) to give an unbiased prior. Then SCOIT updates simultaneously **O, C, G, C**′, and **G**′ via Adam optimization algorithm^33^. Adam adopts momentum and an adaptive learning rate, which makes it suitable for sparse and noisy data. We implemented the optimization process with the Pytorch package and performed the computation on the GPU (Nvidia Tesla T4 16G) for acceleration. We summarize the hyperparameters used in all experiments in Supplementary Table S4. We also investigate how hyperparameters affect performance, and the results show that SCOIT is robust to the choice of embedding dimensions within a certain range (Figure S14).

### Cell clustering and graph construction

In our experiment, we applied the *n* × *k* global cell embedding matrix as low-dimensional embeddings for the *n* cells. We then performed the downstream analysis with the embeddings, including cell clustering and NN graph construction.

For the data from sc-GEM, PEA/STA, CITE-seq, and SCoPE2, we conducted *k*-means clustering for the cell embeddings, with *k* set to the number of cell types and other parameters set to the default. For the data from sci-CAR and SNARE-seq, we constructed the KNN graph and SNN graph with the cell embeddings. For the KNN graph, we connected each cell with the *k*-nearest cells, measured by Euclidean distance. For the SNN graph, we calculated the similarity between two cells using the number of shared *k*-nearest neighbors with the Jaccard index (the overlapping neighbors/the union set of neighbors). *k* was set to 20 for the construction of the KNN and SNN graphs. Then we applied Leiden^26^ to perform community detection for the graphs and labeled the cells.

### Benchmark methods

We applied PCA, TSNE^34^, Seurat^25^, scAI^17^, totalVI^20^, Unioncom^22^, and Matcher^21^ to embed cells in a common latent space as benchmark methods. For PCA and TSNE, we concatenated the multiple data matrices and performed the dimensional reduction along the gene dimension to generate low-dimensional embeddings. We applied the functions implemented in the sci-kit-learn package with the default parameters. For Seurat, following the recommended steps, we applied UMAP^35^ to the constructed weighted nearest neighbor graphs to obtain cell embeddings. scAI is designed to combine transcriptomic and epigenomic data, so we excluded it from the data from PEA/STA2, CITE-seq, and SCoPE2. We set the rank of the inferred factor as the number of cell types and other parameters as default. totalVI is designed for CITE-seq data analysis. We applied it to the CITE-seq sequencing data with default settings. Unioncom and Matcher generate a low-dimensional cell representation for each omics. We applied the default settings and concatenated the multiple embeddings as the final output. Then we applied the above clustering and community detection methods for the cell embeddings generated by the benchmark methods.

With regard to imputation, we applied KNNImputer and SimpleImputer in the sci-kit-learn package as benchmark methods. KNNImputer replaces the missing value to the mean value from *k*-nearest neighbors. We used the default settings (*k* = 5). SimpleImputer imputes the missing value with the mean value of the column.

### Evaluation metrics

All clustering and community detection results are measured using the adjusted Rand index (ARI)^36^, normalized mutual information (NMI)^37^, and Fowlkes-Mallows index (FMI)^38^. Given two sets of clusterings (or communities) *U* (true labels) and *V* (predicted labels) on *n* samples, *U* contains *r* clusters {*U*_1_,*U*_2_, …, *U*_*r*_}, and *V* contains *s* clusters {*V*_1_,*V*_2_, …, *V*_*s*_}. *n*_*i j*_ denotes the number of samples belonging to *U*_*i*_ and *V*_*j*_.

ARI, which is the adjusted version of the Rand index, is defined as

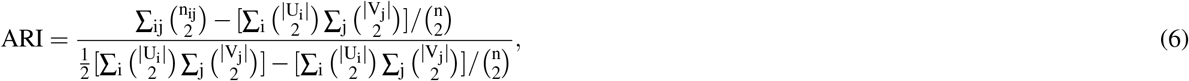

where |*U*_*i*_| and |*V*_*j*_| denote the number of samples in *U*_*i*_ and *V*_*j*_, respectively. ARI gives a similarity score between -1 and 1.

NMI is defined as the mutual information between *U* and *V* divided by the average entropy of *U* and *V*, which can be computed as

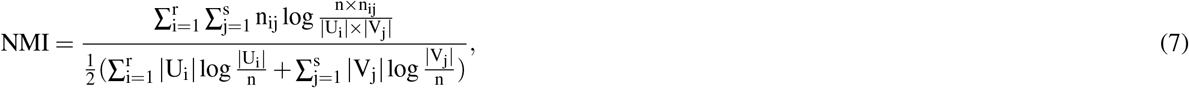

giving a similarity score ranging from 0 to 1.

FMI is defined as the geometric mean between pairwise precision and recall, which is written as

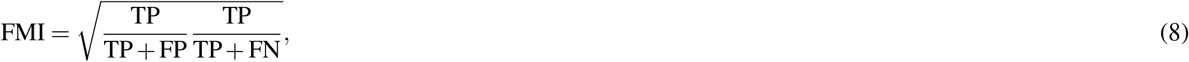

where TP is the number of pairs of samples in the same cluster for both *U* and *V* ; FP is the number of pairs of samples in the same cluster for *V*, but not for *U* ; FN is the number of pair of samples in the same cluster for *U*, but not for *V*. FMI gives a score between 0 and 1.

## Acknowledgements

This work was supported by Strategic Interdisciplinary Research Grant (7020005).

## Author contributions statement

RW designed the software package, performed the experiments, and drafted the manuscript. JW supervised this project and revised the manuscript. SCL proposed the topic, designed the methods, supervised this project, and revised the manuscript. All authors have read and approved the final manuscript.

## References

1. Tang, F. et al. mrna-seq whole-transcriptome analysis of a single cell. Nat. methods 6, 377–382 (2009).

2. Bock, C., Farlik, M. & Sheffield, N. C. Multi-omics of single cells: strategies and applications. Trends biotechnology 34, 605–608 (2016).

3. Argelaguet, R., Cuomo, A. S., Stegle, O. & Marioni, J. C. Computational principles and challenges in single-cell data integration. Nat. biotechnology 39, 1202–1215 (2021).

4. Miao, Z., Humphreys, B. D., McMahon, A. P. & Kim, J. Multi-omics integration in the age of million single-cell data. Nat. Rev. Nephrol. 17, 710–724 (2021).

5. Hu, Y. et al. Single cell multi-omics technology: methodology and application. Front. cell developmental biology 6, 28 (2018).

6. Chappell, L., Russell, A. J. & Voet, T. Single-cell (multi) omics technologies. Annu. review genomics human genetics 19, 15–41 (2018).

7. Cao, J. et al. Joint profiling of chromatin accessibility and gene expression in thousands of single cells. Science 361, 1380–1385 (2018).

8. Yan, R. et al. Decoding dynamic epigenetic landscapes in human oocytes using single-cell multi-omics sequencing. Cell Stem Cell 28, 1641–1656 (2021).

9. Luo, C. et al. Single nucleus multi-omics identifies human cortical cell regulatory genome diversity. Cell genomics 2, 100107 (2022).

10. Cheow, L. F. et al. Single-cell multimodal profiling reveals cellular epigenetic heterogeneity. Nat. methods 13, 833–836 (2016).

11. Chen, S., Lake, B. B. & Zhang, K. High-throughput sequencing of the transcriptome and chromatin accessibility in the same cell. Nat. biotechnology 37, 1452–1457 (2019).

12. Darmanis, S. et al. Simultaneous multiplexed measurement of RNA and proteins in single cells. Cell reports 14, 380–389 (2016).

13. Stoeckius, M. et al. Simultaneous epitope and transcriptome measurement in single cells. Nat. methods 14, 865–868 (2017).

14. Specht, H. et al. Single-cell proteomic and transcriptomic analysis of macrophage heterogeneity using scope2. Genome biology 22, 1–27 (2021).

15. Huang, M. et al. SAVER: gene expression recovery for single-cell RNA sequencing. Nat. methods 15, 539–542 (2018).

16. Chen, H. et al. Assessment of computational methods for the analysis of single-cell atac-seq data. Genome biology 20, 1–25 (2019).

17. Jin, S., Zhang, L. & Nie, Q. scai: an unsupervised approach for the integrative analysis of parallel single-cell transcriptomic and epigenomic profiles. Genome biology 21, 1–19 (2020).

18. Petegrosso, R., Li, Z. & Kuang, R. Machine learning and statistical methods for clustering single-cell RNA-sequencing data. Briefings bioinformatics 21, 1209–1223 (2020).

19. Qi, R., Ma, A., Ma, Q. & Zou, Q. Clustering and classification methods for single-cell RNA-sequencing data. Briefings bioinformatics 21, 1196–1208 (2020).

20. Gayoso, A. et al. Joint probabilistic modeling of single-cell multi-omic data with totalvi. Nat. methods 18, 272–282 (2021).

21. Welch, J. D., Hartemink, A. J. & Prins, J. F. Matcher: manifold alignment reveals correspondence between single cell transcriptome and epigenome dynamics. Genome biology 18, 1–19 (2017).

22. Cao, K., Bai, X., Hong, Y. & Wan, L. Unsupervised topological alignment for single-cell multi-omics integration. Bioinformatics 36, i48–i56 (2020).

23. Argelaguet, R. et al. Multi-Omics Factor Analysis—a framework for unsupervised integration of multi-omics data sets. Mol. systems biology 14, e8124 (2018).

24. Argelaguet, R. et al. MOFA+: a statistical framework for comprehensive integration of multi-modal single-cell data. Genome biology 21, 1–17 (2020).

25. Hao, Y. et al. Integrated analysis of multimodal single-cell data. Cell 184, 3573–3587 (2021).

26. Traag, V. A., Waltman, L. & Van Eck, N. J. From Louvain to Leiden: guaranteeing well-connected communities. Sci. reports 9, 1–12 (2019).

27. Han, H. et al. TRRUST v2: an expanded reference database of human and mouse transcriptional regulatory interactions. Nucleic acids research 46, D380–D386 (2018).

28. Xu, J. et al. A hierarchical integration deep flexible neural forest framework for cancer subtype classification by integrating multi-omics data. BMC bioinformatics 20, 1–11 (2019).

29. Xu, H., Gao, L., Huang, M. & Duan, R. A network embedding based method for partial multi-omics integration in cancer subtyping. Methods 192, 67–76 (2021).

30. Iacono, G. et al. bigScale: an analytical framework for big-scale single-cell data. Genome research 28, 878–890 (2018).

31. Hafemeister, C. & Satija, R. Normalization and variance stabilization of single-cell RNA-seq data using regularized negative binomial regression. Genome biology 20, 1–15 (2019).

32. Stuart, T., Srivastava, A., Lareau, C. & Satija, R. Multimodal single-cell chromatin analysis with Signac. BioRxiv (2020).

33. Kingma, D. P. & Ba, J. Adam: A method for stochastic optimization. arXiv preprint arXiv:1412.6980 (2014).

34. Van der Maaten, L. & Hinton, G. Visualizing data using t-SNE. J. machine learning research 9 (2008).

35. McInnes, L., Healy, J. & Melville, J. Umap: Uniform manifold approximation and projection for dimension reduction. arXiv preprint arXiv:1802.03426 (2018).

36. Steinley, D. Properties of the Hubert-Arable Adjusted Rand Index. Psychol. methods 9, 386 (2004).

37. Vinh, N. X., Epps, J. & Bailey, J. Information theoretic measures for clusterings comparison: Variants, properties, normalization and correction for chance. The J. Mach. Learn. Res. 11, 2837–2854 (2010).

38. Fowlkes, E. B. & Mallows, C. L. A method for comparing two hierarchical clusterings. J. Am. statistical association 78, 553–569 (1983).

