## Supplementary materials for "Probabilistic tensor decomposition extracts better latent embeddings from single-cell multiomic data"

### S1 Supplementary methods

#### S1.1 Multiple-feature tensor construction and decomposition

As shown in Figure S12, for the multiomic data with different features (e.g., $n$ cells with $m_1$ genes and $m_2$ locations), we construct the multiomic tensor with the union set of features among omics. To be more precise, we catenate the two matrices along the dimension of the feature. Still, we differentiate them along the dimension of omics. Then we can construct a tensor of shape $2 \times n \times (m_1 + m_2)$. We do not consider the missing values in the optimization process. The decomposition process is the same as that of a single-feature tensor, except that we do not calculate the omics-specific cell embedding matrix and omics-specific feature embedding matrix. Since the features have already been distinguished on the omics dimension, we omit the terms to avoid the unnecessary model complexity and reduce the costed memory.

#### S1.2 Local element-wise product of multiple vectors

For element-wise product of multiple vectors, we propose a local product method to reserve the variations in each vector. To be specific, for three vector $\mathbf{a}, \mathbf{b}, \mathbf{c} \in \mathbb{R}^k$, we divide them into three parts of length $k_1$, $k_2$, $k_3$, respectively, where $k_1 + k_2 + k_3 = k$. Then we set the element in the third part for $\mathbf{a}$, the second part for $\mathbf{b}$, and the first part for $\mathbf{c}$ as one. Then the sum of element-wise product of $\mathbf{a}$, $\mathbf{b}$, and $\mathbf{c}$ can be written as

$$\begin{aligned}\sum_{k'=1}^{k} \mathbf{a}_{k'}\mathbf{b}_{k'}\mathbf{c}_{k'} &= \sum_{k'=1}^{k_1} \mathbf{a}_{k'}\mathbf{b}_{k'}\mathbf{c}_{k'} + \sum_{k'=k_1+1}^{k_1+k_2} \mathbf{a}_{k'}\mathbf{b}_{k'}\mathbf{c}_{k'} + \sum_{k'=k_1+k_2+1}^{k_1+k_2+k_3} \mathbf{a}_{k'}\mathbf{b}_{k'}\mathbf{c}_{k'} \\ &= \sum_{k'=1}^{k_1} \mathbf{a}_{k'}\mathbf{b}_{k'} + \sum_{k'=k_1+1}^{k_1+k_2} \mathbf{a}_{k'}\mathbf{c}_{k'} + \sum_{k'=k_1+k_2+1}^{k_1+k_2+k_3} \mathbf{b}_{k'}\mathbf{c}_{k'} \qquad \text{(S1)}\end{aligned}$$

In this case, we force each vector to obtain information during optimization. In our implementation, we discard the parts with elements set to one, which does not affect the results, to reduce the costed memory.

#### S1.3 Geometrical interpretation for the tensor decomposition formulation

We give the geometrical interpretation for the tensor decomposition formulation in our proposed framework. As shown in Figure S13, we regard a value in the multiomic tensor as a point in a 3D space, with the values for three dimensions are fixed; regard an omics, a cell, and a gene as a plane, with the value for one dimension is fixed; regard an omics-specific cell, an omics-specific cell, and a gene-specific cell (or can be regarded as a cell-specific gene) as a line, with the values for two dimensions are fixed. Then we give the decomposition formulation as

$$\hat{\mathcal{T}}_{XYZ} = \sum_{k'=1}^{k} \mathbf{O}_{hk'}\mathbf{C}_{ik'}\mathbf{G}_{jk'} + \mathbf{C}'_{XY} \cdot \mathbf{G}_Z + \mathbf{G}'_{XZ} \cdot \mathbf{C}_Y + \mathbf{C}''_{YZ} \cdot \mathbf{O}_X, \qquad \text{(S2)}$$

while the first term is the value of the point at which three planes intersect; the second, third, and fourth terms are the values of the point at which a line and a plane intersect.

For the multiomic integration task, a gene-specific cell or a cell-specific gene representation is of less biological meaning and leads to high memory demands since the numbers of cells and genes are large. Therefore, we skip the last term in our implementation.

#### S1.4 Reparamterization of negative binomial distribution

Let $\mathcal{T}_{hij}$ denotes the observation of gene $j$ in cell $i$ for omics $h$. We write the negative bionomial distribution in Gamma–Poisson mixture format.

$$\begin{aligned} \mathcal{T}_{hij}|\lambda &\sim \mathrm{Poisson}(\lambda_{\mathrm{hij}}) \\ \lambda &\sim \mathrm{Gamma}(\mathrm{r_{hij}}, \mathrm{b_{hij}}) \end{aligned} \tag{S3}$$

Then we can reparameterize the negative bionomial distribution in terms of $r_{hij}$ and $b_{hij}$.

$$\mathcal{T}_{hij} \sim \mathrm{NB}(\mathrm{r_{hij}}, \frac{\mathrm{b_{hij}}}{1+\mathrm{b_{hij}}}) \tag{S4}$$

Let $\mu_{hij}$ be the mean and $\upsilon_{hij}$ be the variance of Gamma distribution. We have $r_{hij} = \frac{\mu_{hij}^2}{\upsilon_{hij}}$ and $b_{hij} = \frac{\mu_{hij}}{\upsilon_{hij}}$. The negative bionomial distribution can be expressed as

$$\mathcal{T}_{hij} \sim \mathrm{NB}(\frac{\mu_{\mathrm{hij}}^2}{\upsilon_{\mathrm{hij}}}, \frac{\mu_{\mathrm{hij}}}{\mu_{\mathrm{hij}}+\upsilon_{\mathrm{hij}}}). \tag{S5}$$

Since the Fano factor assumption suppose the variance scales linearly with the mean as $\upsilon_{hij} = \beta_{hj}\mu_{hij}$. We reparamterize the negative bionomial distribution again.

$$\mathcal{T}_{hij} \sim \mathrm{NB}(\frac{\mu_{\mathrm{hij}}}{\beta_{\mathrm{hj}}}, \frac{1}{1+\beta_{\mathrm{hj}}}) \tag{S6}$$

### S2 Supplementary tables and figures

| Transcription factors | Effect | Target gene |
|---|---|---|
| E2F1 | Activation | AR, ATM, BBC3, CDK1, DHFR, ERCC5, HMGA1, KIF2C, MAP3K5, PCNA, RRM2, THBS1, TP53, TP73 |
| | Repression | BRCA1, DIRAS3, DNMT1, HSPA5, KIF2C, TP53 |
| TP53 | Activation | BBC3, CTSD, DDB1, DNMT1, EPHA2, GSTP1, IFI16, IGFBP3, MMP2, PDGFRB, SFN, THBS1, TP53, TP53I3 |
| | Repression | AFP, BRCA1, BRCA2, CDK1, DDB1, DNMT1, E2F1, HSP90AB1 IGF1R, MAP4, MCM7, MMP1, PRC1, REEP5, WRN, XPO1 |

Table S1: **Summary on the transcription factors and their target genes analyzed in the work.**

| ADT signal (clr-transformed) | Cell type |
|---|---|
| cd11c+,cd14+ | Monocytes |
| cd11c+,cd14- | DC |
| cd34+ | HSC |
| cd56+ | NK |
| cd3+,cd4+,cd8+ | CD8 T cell |
| cd3+,cd4+,cd8- | CD4 T cell |
| cd19+ | B cell |

Table S2: **Epitope-based labeling for CITE-seq data.** We applied clr-transformed ADT signals to discern cell types for the cord blood mononuclear cells from CITE-seq. "+" indicates a positive value; "-" indicates a negative value.

| Data type | Sequencing protocol | Sample | Number of cells | Number of features |
|---|---|---|---|---|
| RNA expression and DNA methylation | sc-GEM | Human fibroblast cell | 224 | 34 genes in RNA expression datasets; 27 genes in DNA methylationd datasets |
| RNA expression and ATAC | sci-CAR | Adult mouse kidney | 8,837 | 17,239 genes in RNA expression datasets; 823,813 peaks in ATAC datasets |
| | SNARE-seq | Neonatal mouse cerebral cortices | 5,081 | 19,322 genes in RNA expression datasets; 229,429 peaks in ATAC datasets |
| | SNARE-seq | Adult mouse cerebral cortices | 10,309 | 33,160 genes in RNA expression datasets; 244,544 peaks in ATAC datasets |
| RNA expression and proteomics | PEA/STA | Human glioblastoma cell | 210 | 88 genes in RNA expression datasets; 78 genes in proteomics datasets |
| | CITE-seq | Cord blood mononuclear cells | 8,617 | 36,280 genes in RNA expression datasets; 13 proteins in proteomics datasets |
| | SCoPE2 | Innate immune cells | 1,490 | 2,272 genes in RNA expression datasets; 3,042 genes in proteomics datasets |

Table S3: **Summary of single-cell multiomic datasets used in this study.**

| Data | Parameter settings |
|---|---|
| Human fibroblast cell (sc-GEM) | $k_1, k_2, k_3$=30, dist="negative_bionomial", lr=1e-2, opt="Adam", n_epochs=260, batch_size=256, $\lambda_c$=0.01, $\lambda_g$=0.01, $\lambda_o$=0.01, $\lambda_{c'}$=1, $\lambda_{g'}$=1 |
| Adult mouse kidney (sci-CAR) | $k_1, k_2, k_3$=20, dist="gaussian", lr=1e-3, opt="Adam", n_epochs=500, batch gradient descent $\lambda_c$=0.01, $\lambda_g$=0.01, $\lambda_o$=0.01 |
| Neonatal mouse cerebral cortices (SNARE-seq) | $k_1, k_2, k_3$=20, dist="gaussian", lr=1e-3, opt="Adam", n_epochs=1700, batch gradient descent $\lambda_c$=0.01, $\lambda_g$=0.01, $\lambda_o$=0.01 |
| Adult mouse cerebral cortices (SNARE-seq) | $k_1, k_2, k_3$=20, dist="gaussian", lr=1e-3, opt="Adam", n_epochs=600, batch gradient descent $\lambda_c$=0.01, $\lambda_g$=0.01, $\lambda_o$=0.01 |
| Human glioblastoma cell (PEA/STA) | $k_1, k_2, k_3$=10, dist="gaussian", lr=1e-2, opt="Adam", n_epochs=850, batch_size=256, $\lambda_c$=0.01, $\lambda_g$=1, $\lambda_o$=1, $\lambda_{c'}$=1, $\lambda_{g'}$=0.01 |
| Cord blood cerebral cortices (CITE-seq) | $k_1, k_2, k_3$=30, dist="gaussian", lr=1e-3, opt="Adam", n_epochs=1500, batch gradient descent $\lambda_c$=0.01, $\lambda_g$=0.01, $\lambda_o$=0.01 |
| Innate immune cells (SCoPE2) | $k_1, k_2, k_3$=30, dist="gaussian", lr=1e-3, opt="Adam", n_epochs=7000, batch gradient descent $\lambda_c$=0.01, $\lambda_g$=0.01, $\lambda_o$=0.01 |
| *in silico* sc-GEM dataset | $k_1, k_2, k_3$=30, dist="negative_bionomial", lr=1e-2, opt="Adam", n_epochs=1000, batch_size=256, $\lambda_c$=0.01, $\lambda_g$=0.01, $\lambda_o$=0.01, $\lambda_{c'}$=1, $\lambda_{g'}$=1 |

Table S4: **Summary of hyperparameters used in this study.**

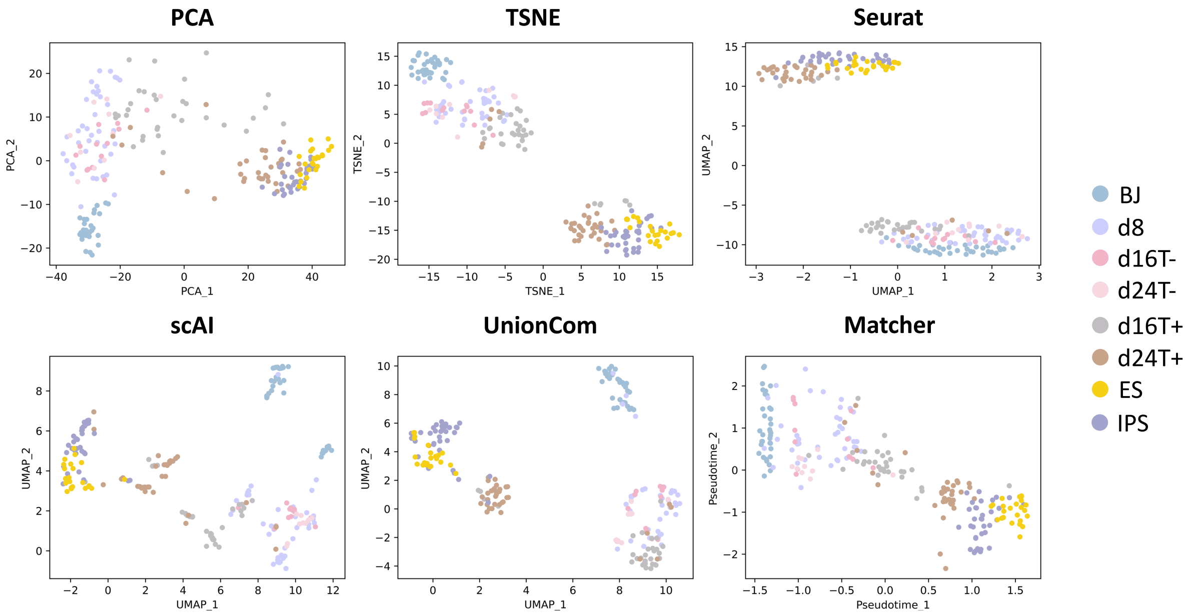


Figure S1: **The UMAP projections of the cell embeddings generated by the benchmark methods for sc-GEM sequencing data.** For human fibroblast cells sequenced with sc-GEM, we show UMAP projections of the cell embeddings generated by PCA, TSNE, Seurat, scAI, UnionCom, and Matcher. Each point represents a cell, color-coded by the true label. The cell types are shown on the right of the scatter plots.

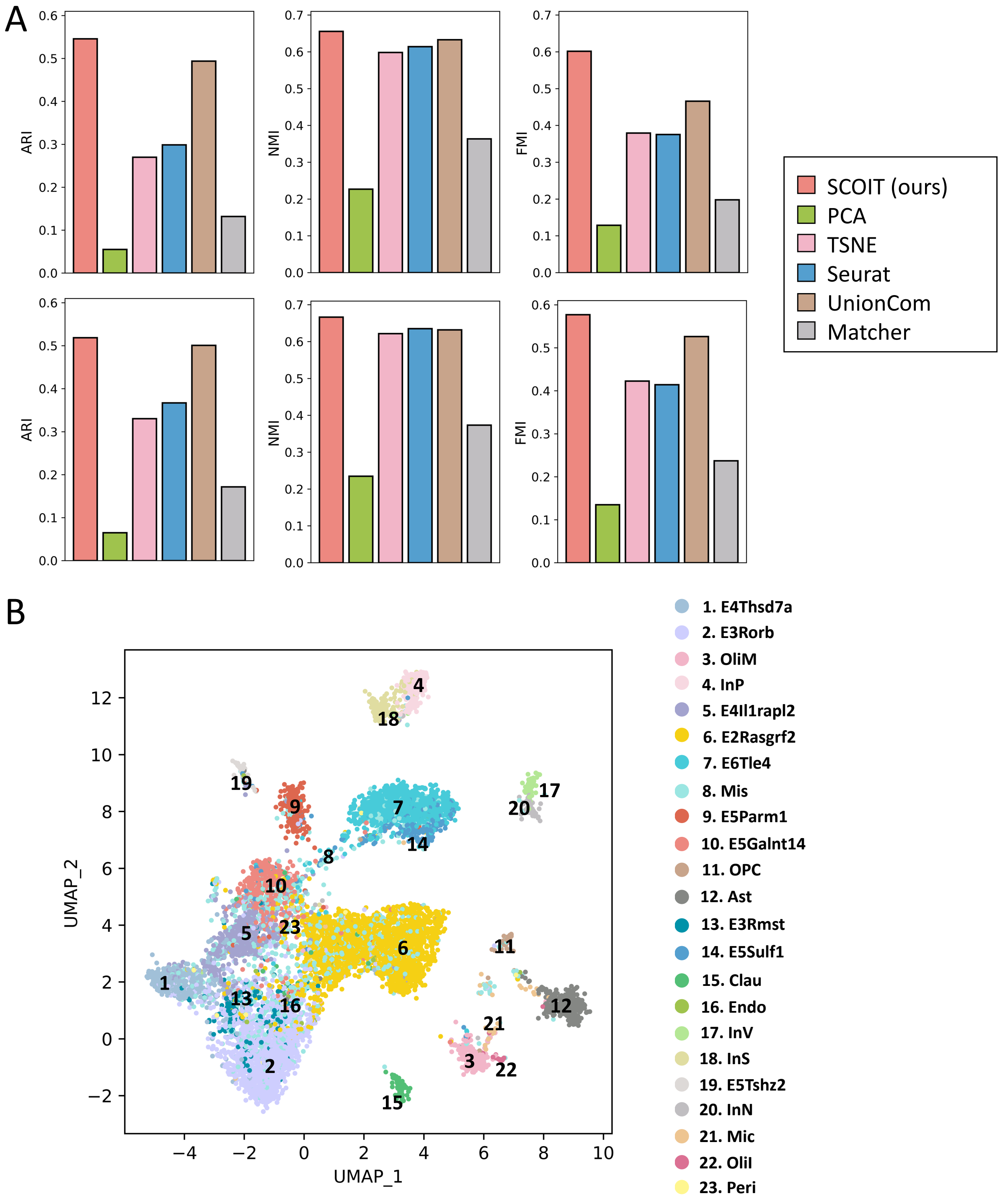


Figure S2: **SCOIT reveals cellular heterogeneity from integrating RNA expression and chromatin accessibility data of adult mouse cerebral cortices cells sequenced with SNARE-seq.** **A.** We compare the Leiden community detection performance for KNN graph (top) and SNN graph (bottom), constructed with cell embeddings generated by different methods. scAI fails to generate results within the time limit (two days). The performance is measured by ARI, NMI, and FMI. **B.** We show the UMAP projections of the global cell embeddings generated by SCOIT. Each point represents a cell, color-coded by the true label. The cell types are shown on the right of the scatter plots.

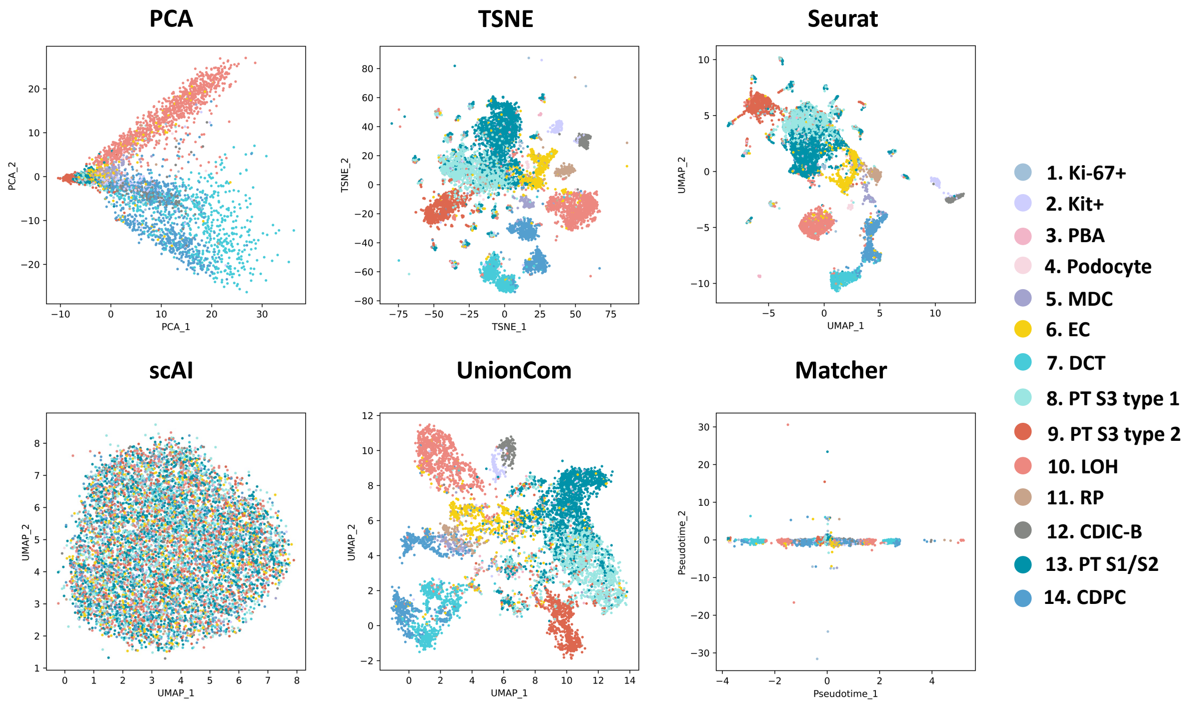


Figure S3: **The UMAP projections of the cell embeddings generated by the benchmark methods for sci-CAR sequencing data.** For adult mouse kidney cells sequenced with sci-CAR, we show UMAP projections of the cell embeddings generated by PCA, TSNE, Seurat, scAI, UnionCom, and Matcher. Each point represents a cell, color-coded by the true label. The cell types are shown on the right of the scatter plots.

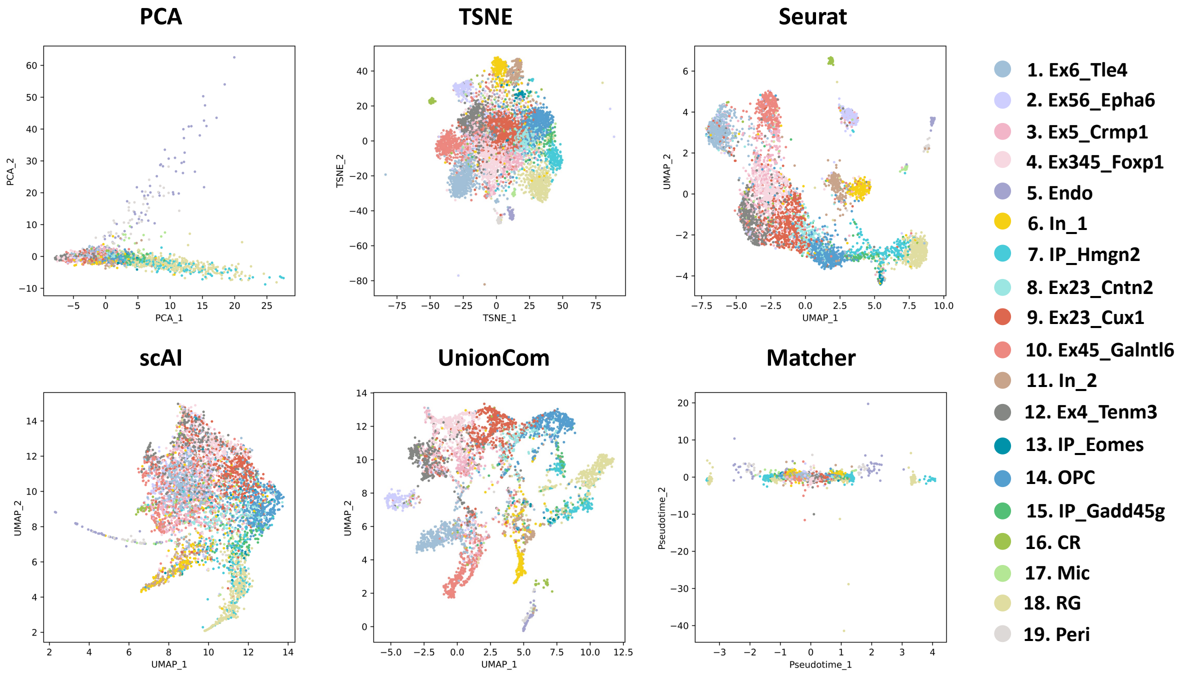


Figure S4: **The UMAP projections of the cell embeddings generated by the benchmark methods for SNARE-seq sequencing data (neonatal mouse cerebral cortices cells).** For neonatal mouse cerebral cortices cells sequenced with SNARE-seq, we show UMAP projections of the cell embeddings generated by PCA, TSNE, Seurat, scAI, UnionCom, and Matcher. Each point represents a cell, color-coded by the true label. The cell types are shown on the right of the scatter plot.

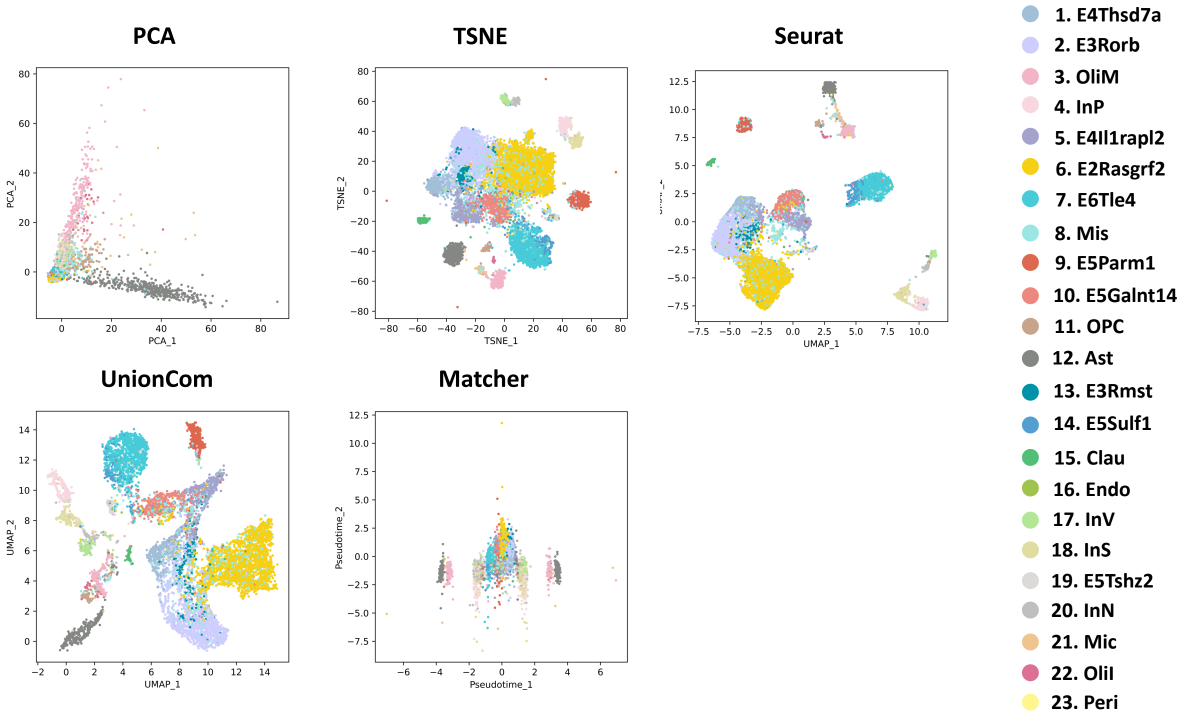


Figure S5: **The UMAP projections of the cell embeddings generated by the benchmark methods for SNARE-seq sequencing data (adult mouse cerebral cortices cells).** For adult mouse cerebral cortices cells sequenced with SNARE-seq, we show UMAP projections of the cell embeddings generated by PCA, TSNE, Seurat, UnionCom, and Matcher. Each point represents a cell, color-coded by the true label. The cell types are shown on the right of the scatter plots.

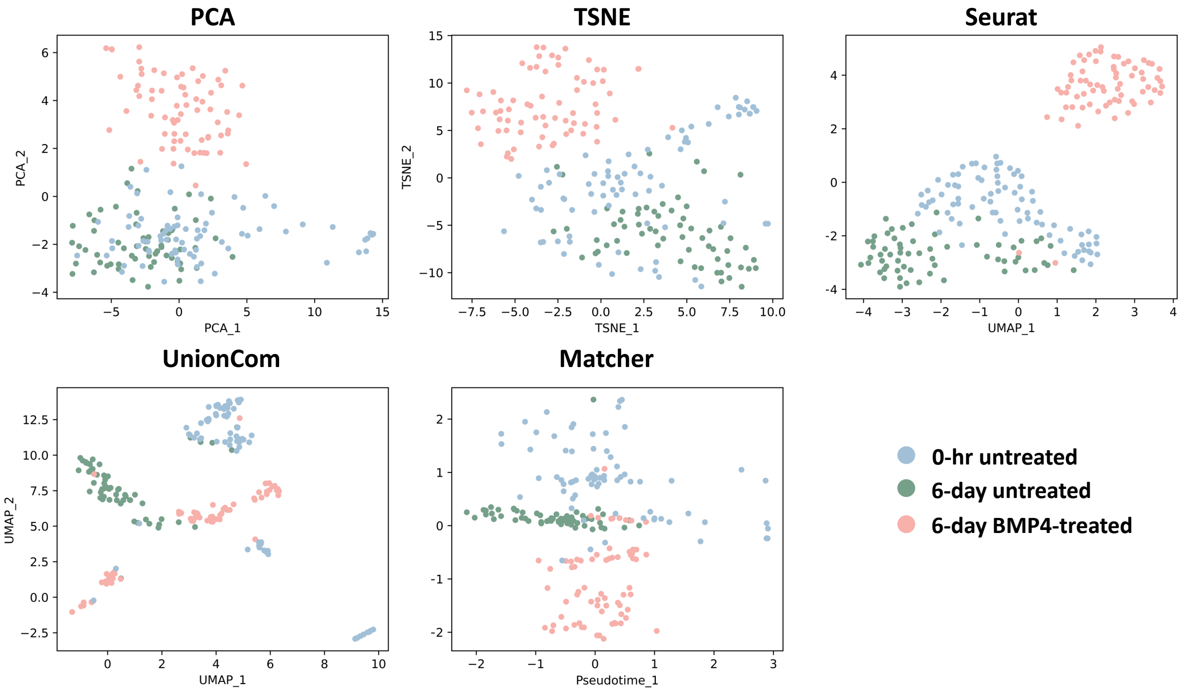


Figure S6: **The UMAP projections of the cell embeddings generated by the benchmark methods for PEA/STA sequencing data.** For human breast adenocarcinoma cells sequenced with PEA/STA, we show UMAP projections of the cell embeddings generated by PCA, TSNE, Seurat, UnionCom, and Matcher. Each point represents a cell, color-coded by the true label. The cell types are shown on the right of the scatter plots.

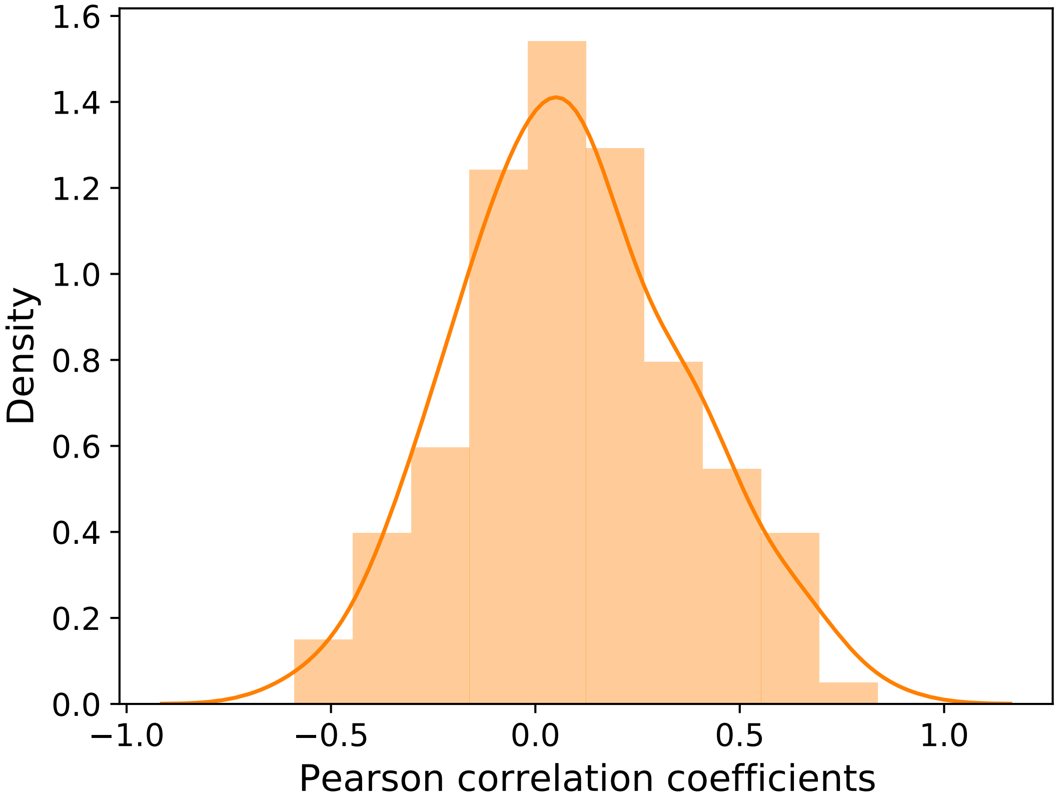


Figure S7: **The distribution of the correlation between gene embeddings from RNA expression and proteomics for PEA/STA sequencing data, generated by SCOIT.** For human breast adenocarcinoma cells sequenced with PEA/STA, we calculate the Pearson correlation coefficients for the omics-specific gene embeddings. The orange line shows the correlation distribution, centered at around zero. The observation indicates less correlation between RNA and protein expression levels for the most gene.

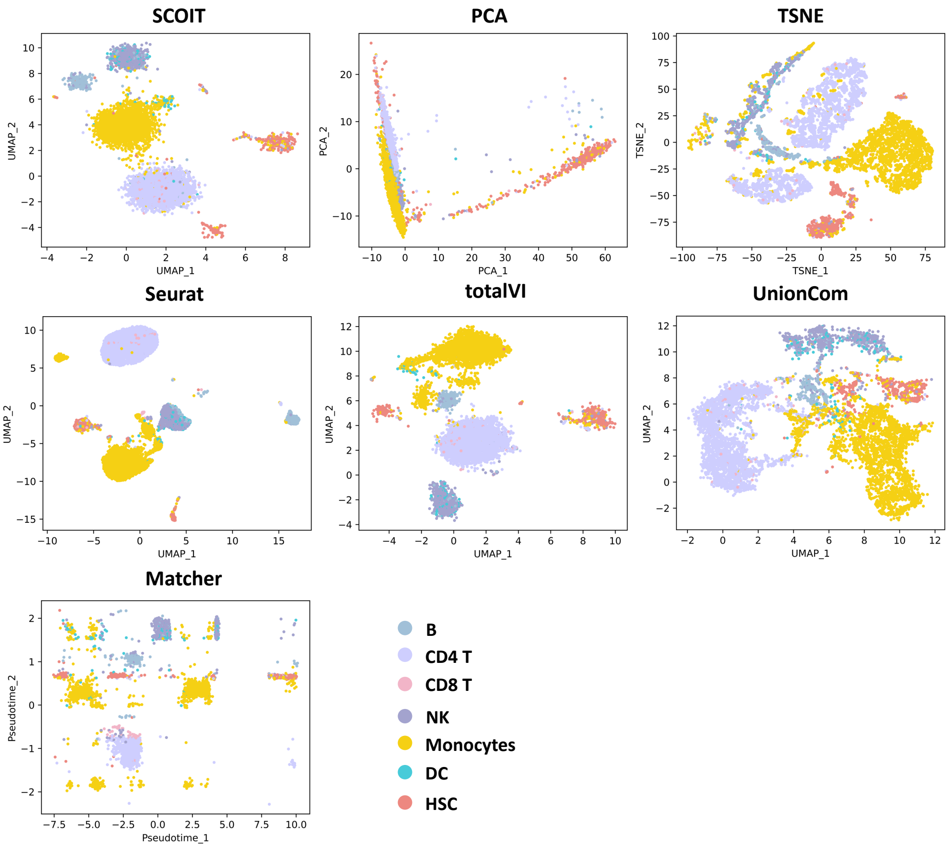


Figure S8: **The UMAP projections of the cell embeddings generated by the different methods for CITE-seq sequencing data.** For cord blood mononuclear cells sequenced with CITE-seq, we show UMAP projections of the cell embeddings generated by SCOIT (our methods), PCA, TSNE, Seurat, totalVI, UnionCom, and Matcher. Each point represents a cell, color-coded by the true label. The cell types are shown on the bottom-right of the panel.

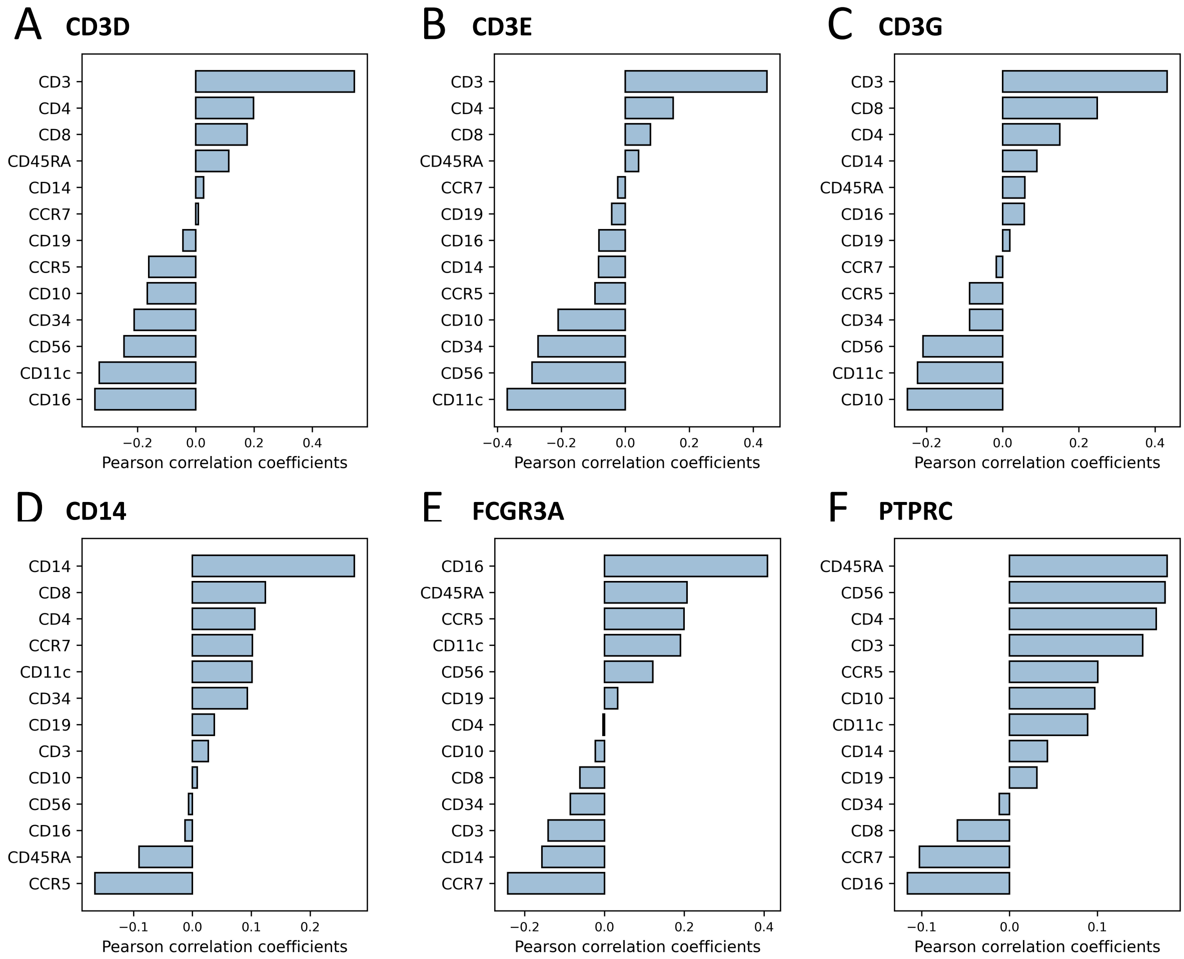


Figure S9: **The correlation between gene embeddings and epitope embeddings generated by SCOIT for CITE-seq sequencing data.** We choose six genes (CD3D, CD3E, CD3G, CD14, FCGR3A, and PTPRC) that encode the corresponding epitope. Then we calculate the correlation between the gene embeddings and the 13 epitope embeddings generated by SCOIT. The epitopes are sorted by the Pearson correlation coefficients.

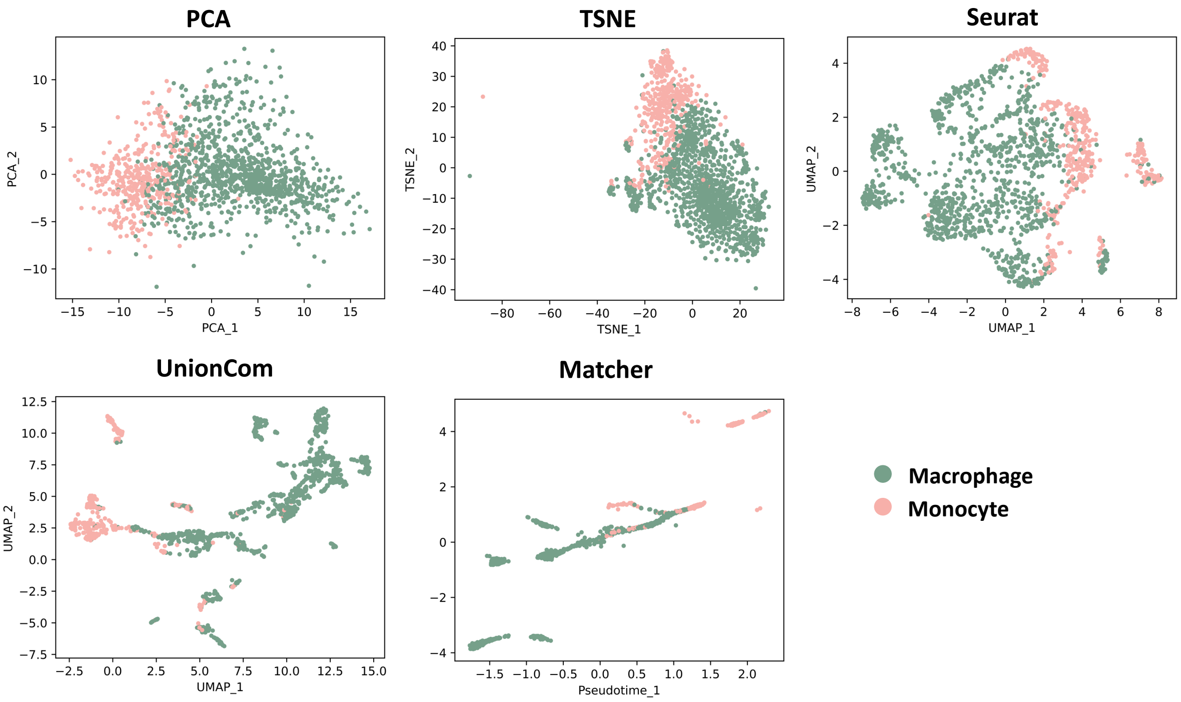


Figure S10: **The UMAP projections of the cell embeddings generated by the different methods for SCoPE2 sequencing data.** For innate immune cells sequenced with SCoPE2, we show UMAP projections of the cell embeddings generated by PCA, TSNE, Seurat, UnionCom, and Matcher. Each point represents a cell, color-coded by the true label. The cell types are shown on the bottom-right of the panel.

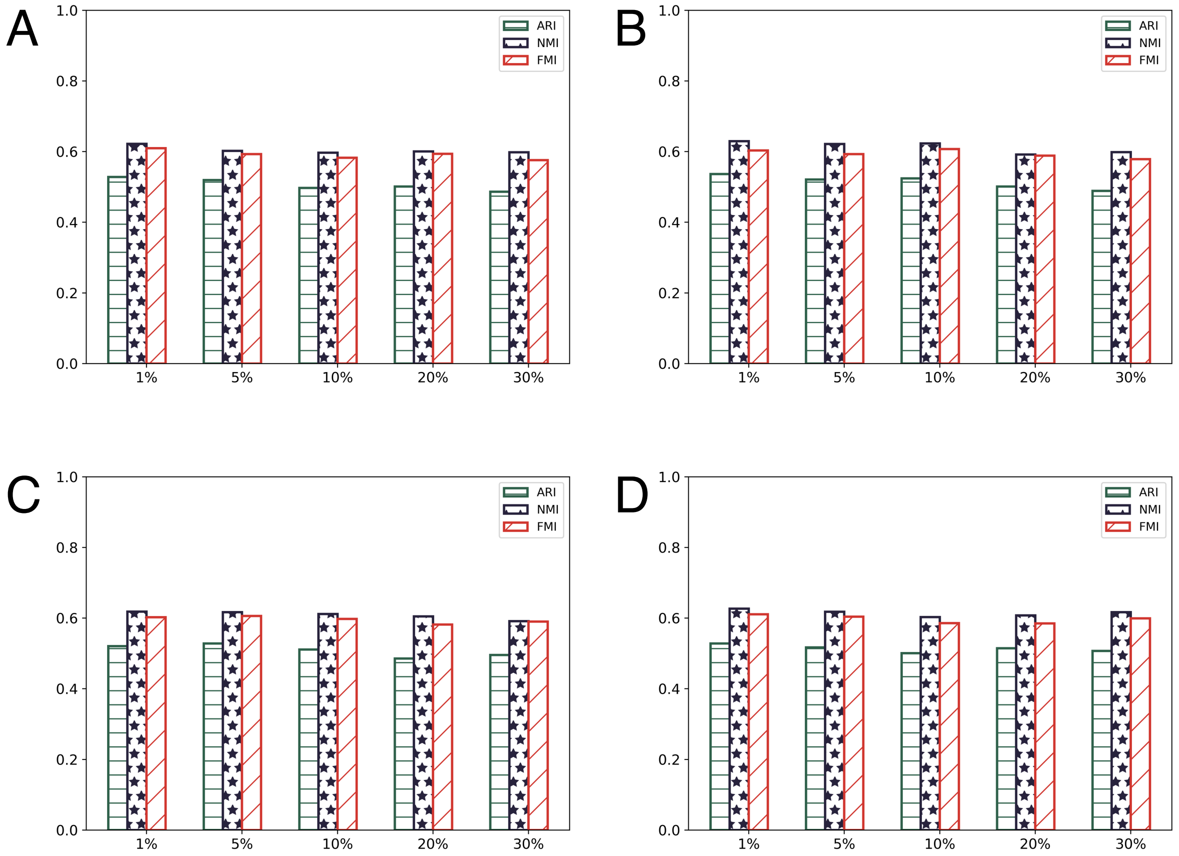


Figure S11: **The $k$-means clustering performance on SCOIT generated global cell embeddings.** We add missing values and Gaussian noise to the sc-GEM dataset to assess SCOIT for sparse and noisy data. **A**. We set $1\% - 30\%$ of the RNA expression data as the missing value. **B**. We set $1\% - 30\%$ of the DNA methylation data as the missing value. **C**. We add Gaussian noise to $1\% - 30\%$ of the RNA expression data. **D**. We add Gaussian noise to $1\% - 30\%$ of the DNA methylation data. The performances are measured by ARI, NMI, and FMI. The higher bar corresponds to more concordance between the predicted and true labels.

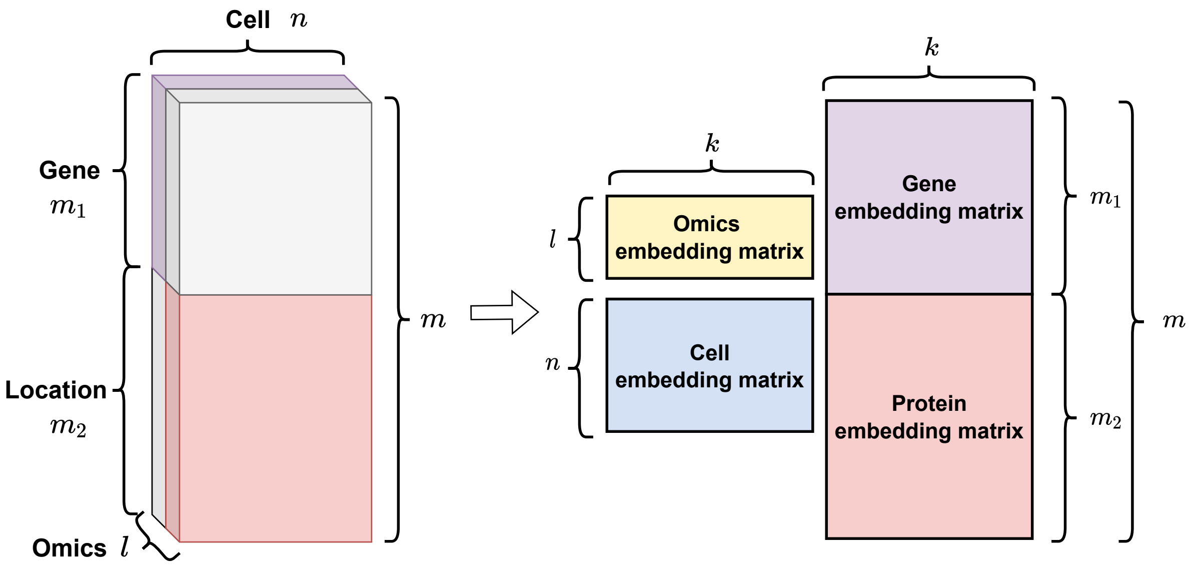


Figure S12: **The construction and decomposition of a multiple-feature tensor.** We combine a data matrix of shape $n \times m_1$ (purple) and a data matrix of shape $n \times m_2$ (pink) into a tensor of shape $2 \times n \times m$, while $m = m_1 + m_2$. The gray parts can be regarded as missing values. We decompose the tensor into a omics embedding matrix ($l \times k$), a cell embedding matrix ($n \times k$), a gene embedding matrix ($m_1 \times k$) and a protein embedding matrix ($m_2 \times k$).

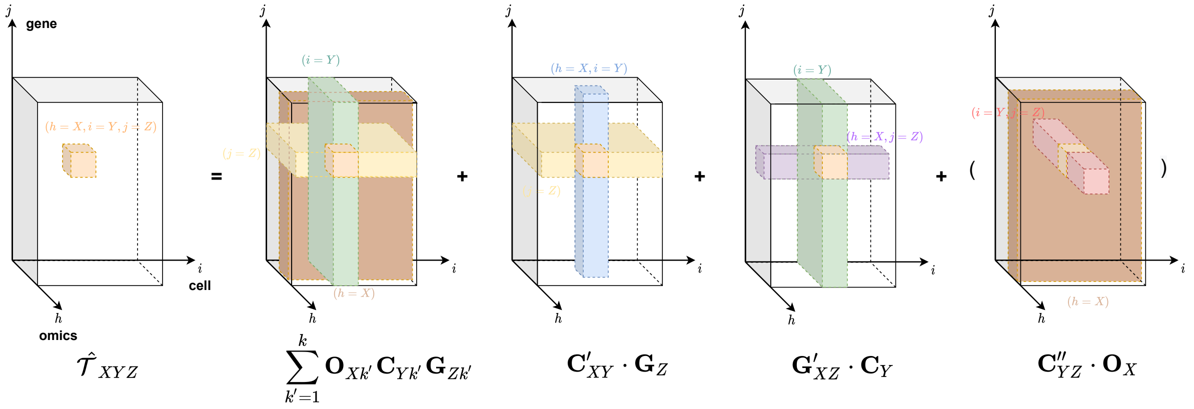


Figure S13: **The geometrical interpretation for the tensor decomposition formulation.** The orange point represents a value in a third-order multiomic tensor, denoted as $\hat{\mathcal{T}}_{XYZ}$; The brown plane represents an omics, denoted as $\mathbf{O}_X$; The green plane represents a cell, denoted as $\mathbf{C}_Y$; The yellow plane represents a gene, denoted as $\mathbf{G}_Z$; The blue line represents an omics-specific cell, denoted as $\mathbf{C}'_{XY}$; The purple line represents an omics-specific gene, denoted as $\mathbf{G}'_{XZ}$; The pink line represents a gene-specific cell (or can be regarded as a cell-specific gene), denoted as $\mathbf{C}''_{YZ}$ (or $\mathbf{G}''_{YZ}$).

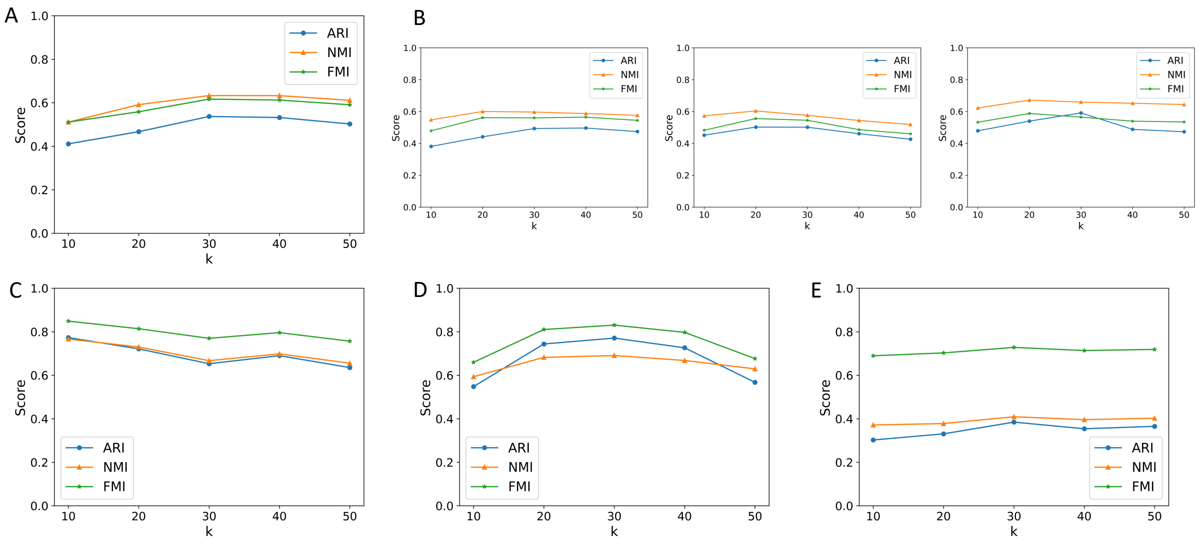


Figure S14: **Clustering performance assessment with regard to the embedding dimension.** For **A**. human fibroblast cells from sc-GEM, **B**. adult mouse kidney cells from sci-CAR, neonatal mouse cerebral cortices cells and adult mouse cerebral cortices cells from SNARE-seq (from left to right), **C**. human glioblastoma cells from PEA/STA, **D**. cord blood cerebral cortices cells from CITE-seq, and **E**. innate immune cells from SCoPE2, we show the clustering performance of SCOIT is robust to the choice of $k_1$, $k_2$, and $k_3$ within certain range. Further increasing the embedding dimensions leads to overfitting.
